# A naturalistic reinforcement learning task uncovers historical neural representations

**DOI:** 10.1101/2025.07.17.665306

**Authors:** Sangsoo Jin, Juhyeon Lee, Satrajit Ghosh, Jong-Hwan Lee

**Affiliations:** Department of Brain and Cognitive Engineering, Korea University, Seoul, Republic of Korea; McGovern Institute for Brain Research, MIT, Cambridge, MA 02139, USA; Department of Otolaryngology Head and Neck Surgery, Harvard Medical School, Boston, MA, USA; Interdisciplinary Program in Precision Public Health, Korea University, Seoul, Republic of Korea

**Keywords:** Functional magnetic resonance imaging, Historical learning, Naturalistic paradigm, Reinforcement learning

## Abstract

Exploration is essential for reinforcement learning (RL) in human development, facilitating cognitive and behavioral adaptation. However, conventional RL paradigms in neuroimaging studies often rely on highly constrained search spaces and artificial, non-naturalistic scenarios, limiting their capacity to reflect the complexity of real-world human exploration and decision-making. To address this gap, we developed a novel, naturalistic RL paradigm called *Photographer*, which involves a photograph-taking task set in a virtual street-view environment. Participants navigated city scenes and attempted to infer the paradigm’s covert goal solely from feedback, defined as the conceptual similarity between their photograph and a hidden target sentence. Representational similarity analysis revealed that feedback history, an encoding of recent trial attempts, was robustly represented in the middle orbital gyrus and inferior frontal gyrus. These neural representations were consistently observed across two independent participant groups. Moreover, although the two regions coactivated during exploration, each was associated with a distinct functional network.

## Introduction

Reinforcement learning (RL) plays a critical role in human development, guiding behavior based solely on prior positive and negative experiences ^1^. Exploration is essential to effective RL, as it enables agents to identify rewarding and aversive stimuli and transition toward higher-value outcomes. Children and adolescents tend to explore more stochastically than older adults ^2–4^, whereas adults more efficiently adapt their behavior through strategic exploration ^1,5,6^. Reflecting this developmental shift, recent work has proposed that human learning resembles stochastic optimization ^7^, wherein learning strategies begin with high stochasticity and are gradually fine-tuned over the course of development.

Despite the significance of RL in development, most prior neuroimaging studies employing RL paradigms have been conducted in controlled laboratory environments. These studies typically involved tasks such as multi-armed bandits ^8–10^, card deck selections ^1,11,12^, or abstract visual stimuli ^13–15^. While such paradigms introduce stochasticity and dynamic reward contingencies, adult participants tend to adapt rapidly due to the limited and simplified search spaces, which differ markedly from those encountered in real-world RL scenarios. Recent approaches have employed video games to introduce more complex goals and environments ^16,17^, but these designs still fall short of capturing the nuance and ecological validity of real-world learning experiences ^18^, and participants’ prior familiarity with game mechanics may further confound results.

To address these limitations, we developed a novel RL paradigm, called *Photographer*, that simulates a real-world photograph-taking task in a naturalistic environment (Fig. 1a). Using an MR-compatible joystick, participants were afforded a high degree of exploratory freedom, including the ability to look around and navigate between locations, mimicking real-world behavior. Participants received minimal task instructions and were expected to build an internal model of the task environment based solely on visual feedback, thereby engaging in strategic exploration necessary for successful learning. This design aimed to preserve ecological realism while eliciting naturalistic RL behavior. We hypothesized that adult participants would exhibit effective learning behavior in this paradigm, reflected in improved performance based on feedback scores. Additionally, we expected consistent identification of the neural correlates of learning across two independent cohorts (discovery and validation groups), thereby supporting the robustness and reproducibility of the *Photographer* paradigm for neuroimaging research.

**Fig. 1.**
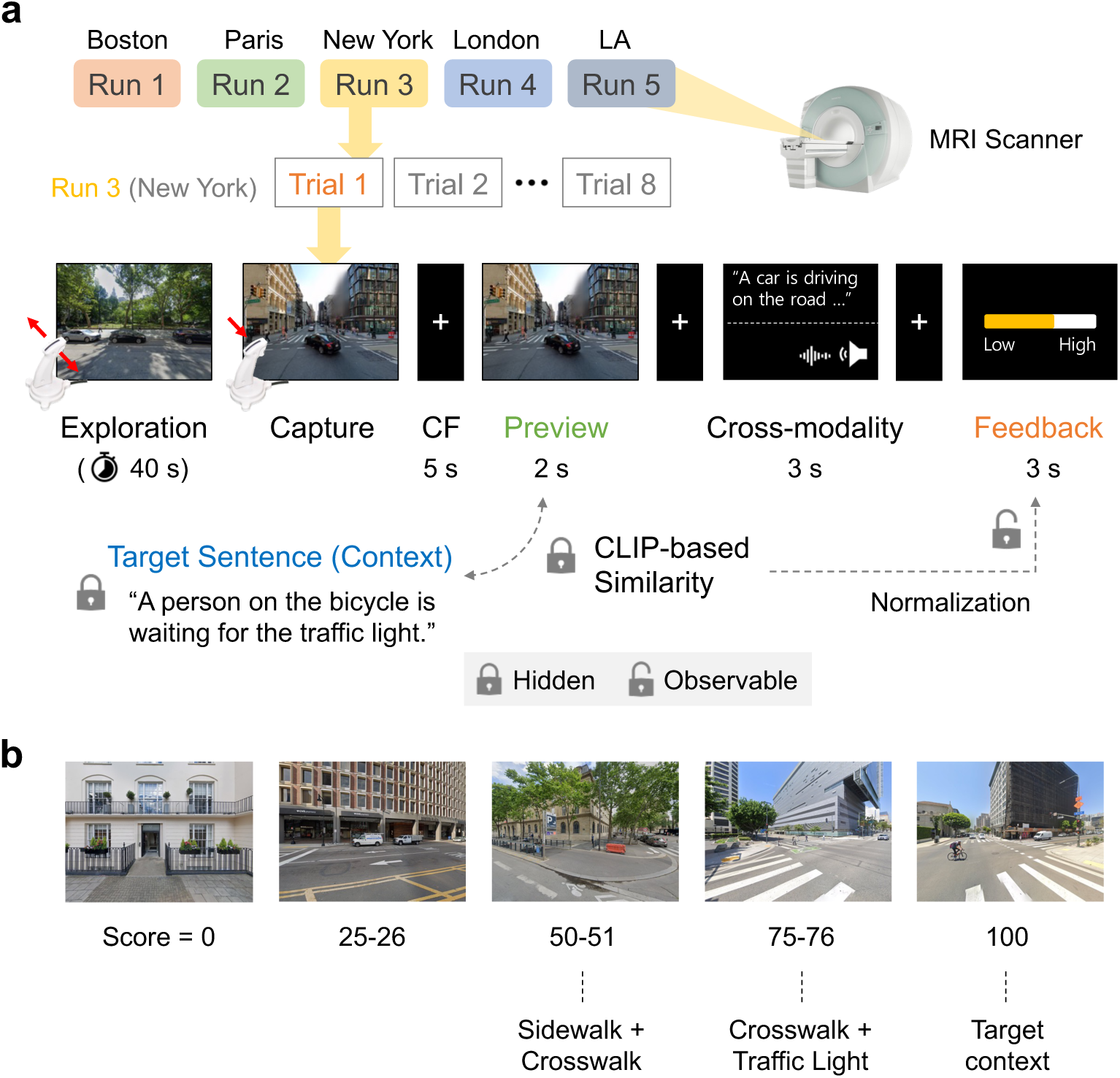
The Photographer paradigm. **a**, Schematic of a single task run. Participants completed five runs, each in a different city visited in pseudorandom order. In each run, they completed eight trials (“photograph attempts”). During each trial, participants explored a Google Street View scene using a joystick (“Exploration”), captured an image (“Capture”), briefly reviewed the photo (“Preview”), and then received a cross-modal image caption generated by a deep learning model, presented either in caption or voice (“Cross-modality”). Finally, a feedback score was displayed (“Feedback”), reflecting the CLIP-based similarity between the participant’s photograph and a hidden target sentence. Participants were instructed to take photos that would yield higher feedback scores, without being informed of the target sentence or the underlying feedback mechanism. This setup allowed participants to implicitly learn the task structure, as only the normalized score was observable. CF, cross-fixation; CLIP, Contrastive Language–Image Pretraining. **b**, Example photographs representing five feedback score ranges (0, [25–26], [50–51], [75–76], and 100). Crosswalks begin to appear at moderate scores (e.g., 25), with high-scoring images depicting target-relevant objects such as traffic lights. The image receiving the highest score closely resembles the hidden target context.

## Results

### Photographs associated with the target sentence yielded higher feedback scores

Photographs (i.e., captured images) containing crosswalks tended to receive higher feedback scores, and prominent scene elements, such as traffic lights and bicycles, began to appear when feedback scores exceeded 75 (Fig. 1b). The presence of specific target objects in the images was associated with significantly increased feedback scores (Fig. 2a). In particular, scenes that included a person yielded significantly higher scores, *t*(442.54) = 4.50, Bonferroni-corrected *P* < 0.001, *d* = 0.30, 95% confidence interval (CI): [0.17, 0.44]; those with a bicycle, *t*(282.10) = 10.45, *P* < 0.001, *d* = 0.74, 95% CI: [0.58, 0.89]; and those with a traffic light, *t*(1261.67) = 12.47, *P* < 0.001, *d* = 0.68, 95% CI: [0.57, 0.88].

**Fig. 2.**
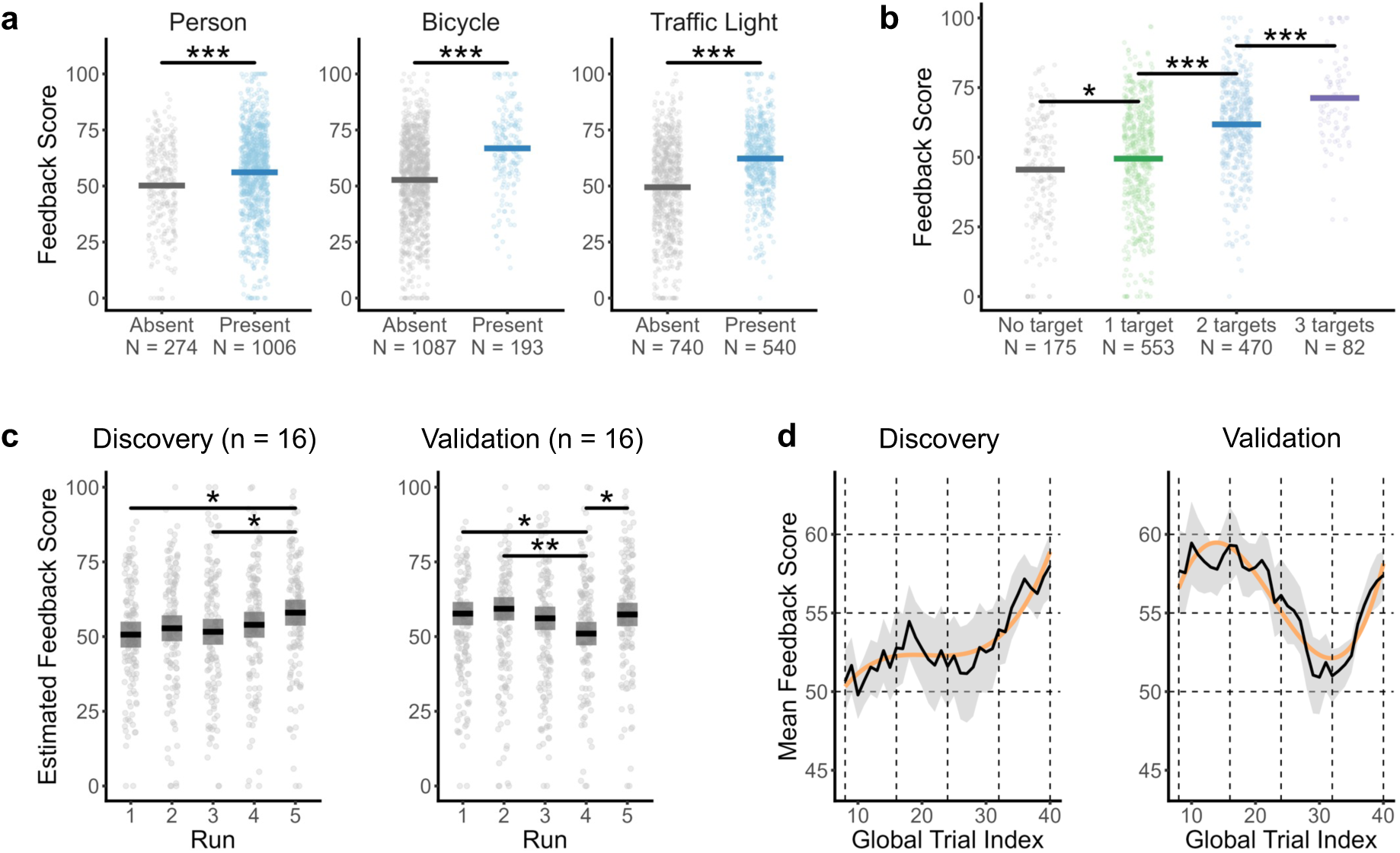
Validation of feedback scores and participant behavior. **a**, Feedback score differences based on the presence or absence of target objects. Two-sample independent *t*-tests were conducted to compare the mean feedback scores of photographs depending on whether each of the three target objects was visible. Thick horizontal lines indicate the mean scores. **b**, Mean feedback scores as a function of the number of target objects detected in the photographs. Scores increased monotonically with the number of target objects. All post hoc pairwise *t*-tests were significant after FDR correction; only three comparisons are annotated for clarity. **c**, Estimated marginal means of feedback scores across the five task runs, shown separately for each subgroup. Dark lines indicate estimated marginal means, and shaded areas represent 95% confidence intervals. Significant pairwise differences between task runs (two-sided *P* < 0.05, FDR- corrected) are annotated. **d**, Group-level moving average trajectories of feedback scores (window size = 8 trials). Individual scores from trials 8 to 40 were aggregated to generate group-level learning curves. Black lines represent group means; shaded regions indicate standard errors; orange curves show linear mixed- effects model estimates. *: *P* < 0.05; **: *P* < 0.01; ***: *P* < 0.001.

Furthermore, feedback scores increased systematically with the number of target objects present in each photograph (Fig. 2b). A significant main effect of object count was observed, *F*(3, 1276) = 76.39, *P* < 0.001, η² = 0.15, one-sided 95% CI: [0.12, 1.00]. All post hoc pairwise *t*-tests yielded FDR-corrected *P* values below 0.05.

### Participants showed evidence of learning

To evaluate whether participants learned the task goals, we analyzed between-run changes in feedback scores using linear mixed-effects (LME) models for both the discovery and validation groups. In the discovery group, task run had a significant main effect on feedback scores when accounting for random effects of subjects and trials (log-likelihood improvement over the null model: χ²(4) = 10.78, *P* = 0.029; Akaike Information Criterion [AIC] improvement: ΔAIC = –2.8; Fig. 2c). Estimated marginal means indicated that feedback scores in Run 5 were significantly higher than in Run 1 (Run 5 – Run 1 = 7.35; *t*(616) = 3.00; FDR-corrected *P* = 0.028; *d* = 0.37) and Run 3 (Run 5 – Run 3 = 6.40; *t*(616) = 2.61; *P* = 0.046; *d* = 0.32).

Similarly, the validation group showed a distinct but significant effect of task run on feedback scores (χ²(4) = 15.02, *P* = 0.005; ΔAIC = –7.0). In this group, estimated marginal means revealed that scores in Run 4 were significantly lower than those in Run 1 (Run 4 – Run 1 = –6.70; *t*(616) = –2.91; *P* = 0.019; *d* = –0.35), Run 2 (Run 4 – Run 2 = –8.30; *t*(616) = –3.61; *P* = 0.003; *d* = –0.44), and Run 5 (Run 4 – Run 5 = –6.40; *t*(616) = –2.78; *P* = 0.019; *d* = –0.34).

Model comparisons of moving-averaged feedback scores further indicated that LME models incorporating a cubic polynomial term provided the best fit for both groups, as evidenced by significant improvements in log-likelihood and AIC values (discovery group: χ²(3) = 33.55, *P* < 0.001, ΔAIC = –27.5; validation group: χ²(3) = 87.51, *P* < 0.001, ΔAIC = –81.5; Fig. 2d). These results suggest that incorporating trial-wise polynomial terms significantly improved the model fit compared to intercept-only models, indicating evidence of between-trial learning dynamics in both groups.

### Historical feedback representations are encoded in the prefrontal cortex

During the presentation of feedback scores, we observed significant activation in the bilateral occipital cortex, including the superior, middle, and inferior occipital gyri, as well as in the supplementary motor area, right middle and inferior frontal gyri (IFG), precentral gyrus, and anterior insula (Supplementary Fig. 1). However, no clusters survived FDR correction when using a parametric regressor representing trial-wise feedback scores, indicating a lack of consistent encoding of feedback at the single-trial level.

In contrast, models incorporating feedback history across multiple trials (i.e., feedback scores of recent trial attempts) revealed robust and replicable activation patterns across both subgroups, as determined by searchlight representational similarity analysis (RSA). Specifically, the *Recent-2 Trial* model identified clusters in the IFG or ventrolateral prefrontal cortex (vlPFC), and in the middle orbital gyrus (MiOG) or ventromedial prefrontal cortex (vmPFC) (Fig. 3a; Supplementary Table 2). The *Recent-3 Trial* model additionally revealed activation in the ventral striatum (VS), postcentral gyrus (PCG) or inferior parietal lobule (IPL), IFG/vlPFC, MiOG/vmPFC, and anterior cingulate cortex (ACC) (Fig. 3b; Supplementary Table 2). Notably, both multiple-trial models showed overlapping activation in the vmPFC and vlPFC, suggesting consistent encoding of feedback history in these prefrontal regions (Fig. 3c; Supplementary Table 2).

**Fig. 3.**
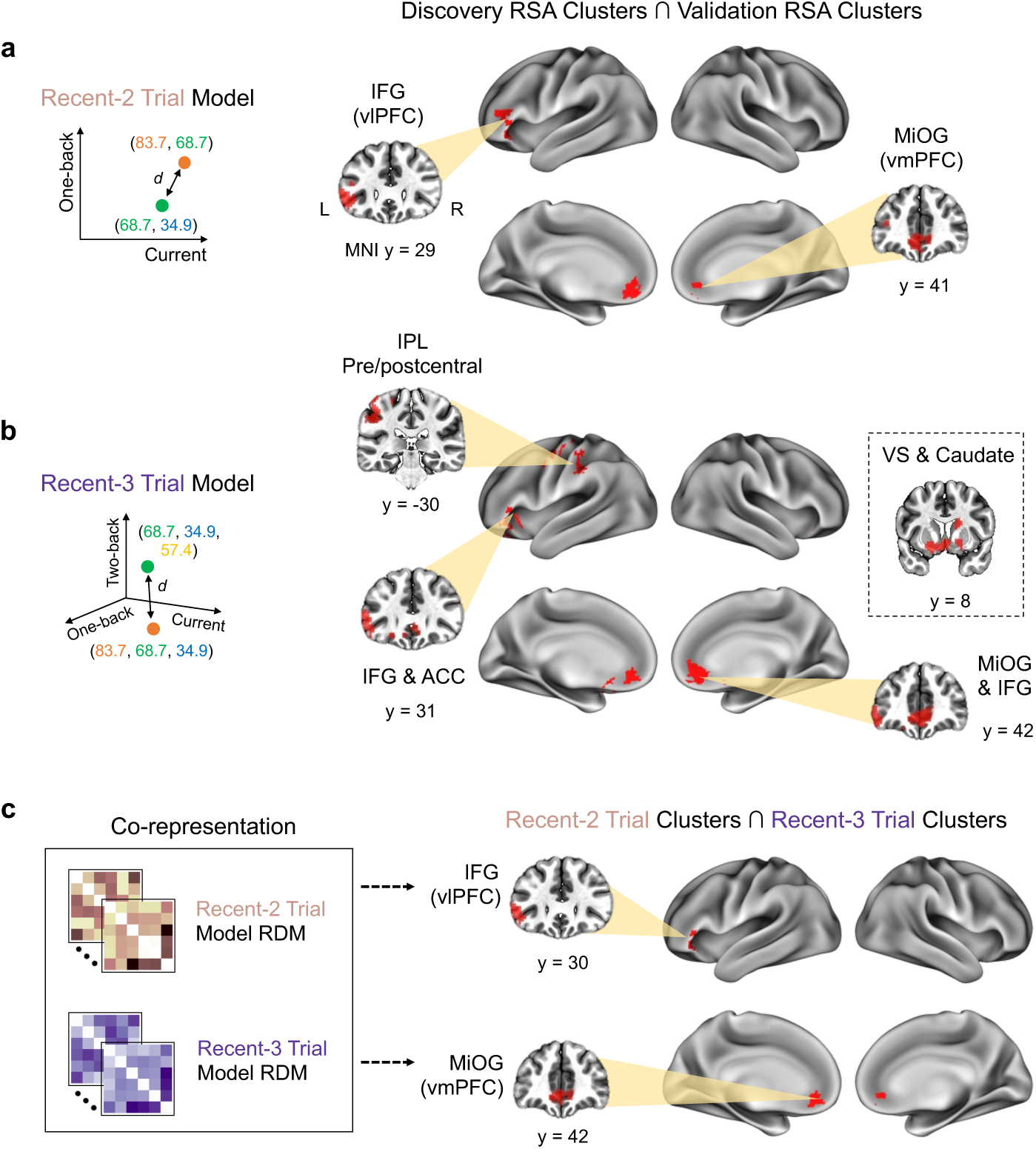
Cross-validated clusters representing feedback history models. Robust representations were observed only for multiple-trial feedback history models across both subgroups. **a**, Clusters significantly encoding the Recent-2 Trial model in both subgroups (voxel-level *P* < 0.005, one-sided; cluster-level α < 0.05; overlapping cluster extent ≥ 20 voxels). **b**, Cross-validated clusters encoding the Recent-3 Trial model. **c**, Clusters jointly representing both the Recent-2 and Recent-3 Trial models across subgroups. All reported clusters contain at least 20 voxels. L, left; R, right; IFG, inferior frontal gyrus; vlPFC, ventrolateral prefrontal cortex; MiOG, middle orbital gyrus; vmPFC, ventromedial prefrontal cortex; IPL, inferior parietal lobule; VS, ventral striatum; ACC, anterior cingulate cortex.

To further validate these findings, we examined three control model representational dissimilarity matrices (RDMs), each encoding feedback history using different combinations of prior trials: the one-back trial, two-back trial (single-trial models), and the *Previous-2 Trial* model (a composite of one- and two-back feedback scores). Of these, only the Previous-2 Trial model yielded cross-validated activation clusters in the MiOG and superior frontal gyrus (SFG) (Supplementary Fig. 2a). Notably, a 24-voxel cluster in the MiOG consistently encoded all three multiple-trial feedback history models, Recent-2 Trial, Recent-3 Trial, and Previous-2 Trial, demonstrating its robust involvement in integrating recent feedback history (Supplementary Fig. 2b).

### The IFG (vlPFC) and MiOG (vmPFC) represent both feedback history and spatial map information

The inferior frontal gyrus (IFG; i.e., ventrolateral prefrontal cortex, vlPFC) and middle orbital gyrus (MiOG; i.e., ventromedial prefrontal cortex, vmPFC) may also encode task-relevant information acquired during exploration, independent of feedback scores presented during the feedback phase. To investigate this possibility, we conducted second-level RSA within the IFG and MiOG clusters identified previously. Specifically, we tested six feedback history model RDMs derived from prior searchlight RSA, alongside three additional behavioral model RDMs encoding exploration-related features.

These three behavioral RDMs captured task behaviors during exploration and capture events: the *exploration time* and *exploration distance* models encoded the temporal duration and spatial extent of each exploration episode, respectively; the *capture distance* model represented geodesic distances between consecutive photograph capture locations, effectively modeling spatial proximity among capture sites. We compared each behavioral model RDM with the neural RDMs extracted from all voxels within the MiOG and IFG cluster regions. For statistical testing, we employed one-sample *t*-tests with 10,000 random permutations per neural-model RDM pairing.

Consistent with previous searchlight RSA results, both MiOG and IFG clusters robustly encoded feedback history models across trials, including the Recent-2, Recent-3, and Previous-2 Trial models (Supplementary Fig. 3). In addition, both regions strongly encoded the capture distance model across the discovery and validation groups (MiOG: discovery, *t*(15) = 9.450, *P* < 0.001, *d* = 2.440; validation, *t*(15) = 6.917, *P* < 0.001, *d* = 1.786; IFG: discovery, *t*(15) = 5.981, *P* < 0.001, *d* = 1.544; validation, *t*(15) = 6.806, *P* < 0.001, *d* = 1.757; Fig. 4a).

**Fig. 4.**
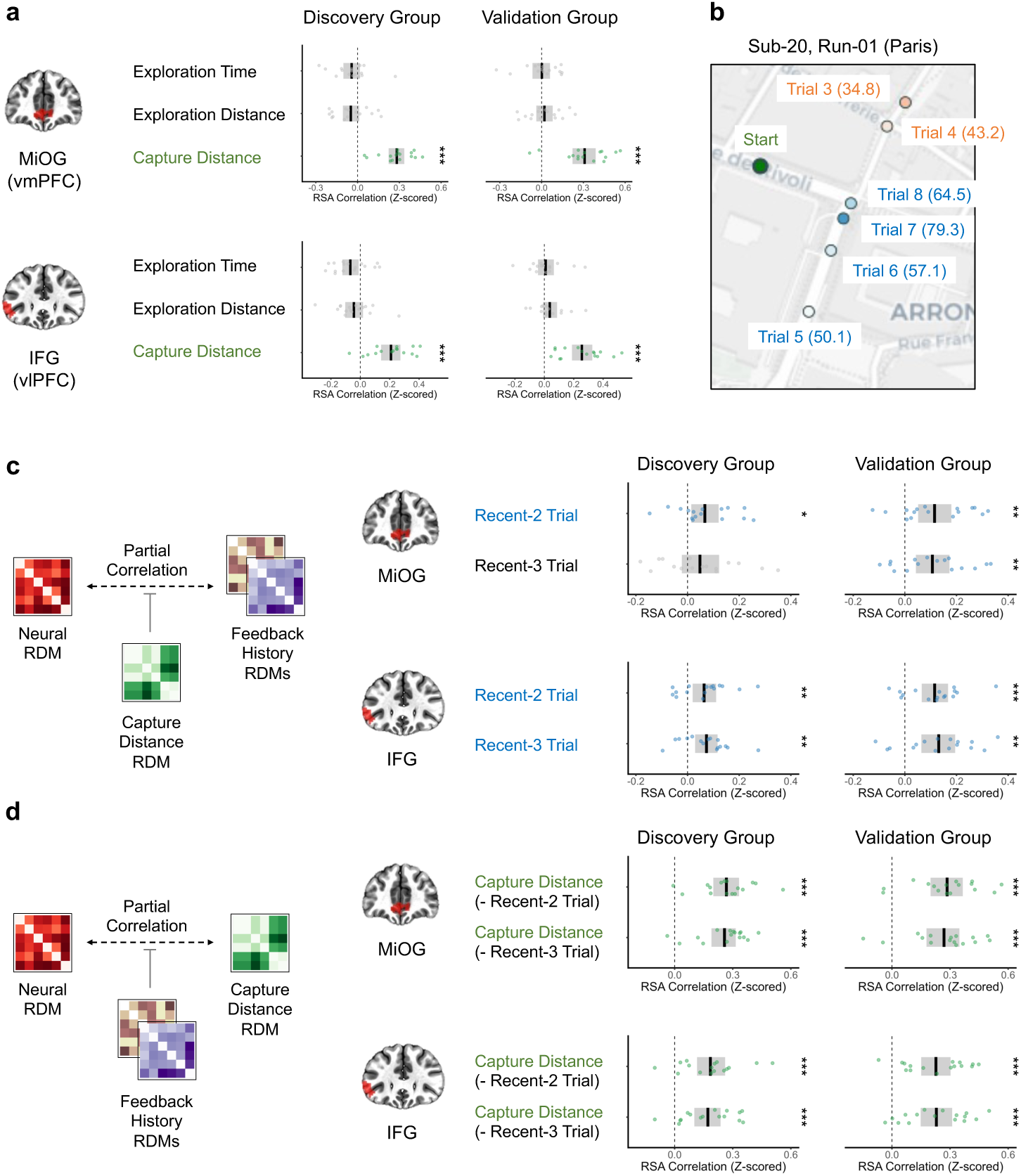
Second-level RSA results for MiOG and IFG clusters. **a**, Group-level representational similarity analysis (RSA) results for exploration-based model RDMs. Neural RDMs were constructed using beta values from the MiOG and IFG masks (defined in Fig. 3c) and compared to three exploration model RDMs. Models showing significant similarity (one-sided *P* < 0.05, based on 10,000 permutation-derived null distributions) are highlighted in green. **b**, Example visualization illustrating the association between spatial adjacency and feedback score similarity across trials. **c**, Partial correlation RSA results between neural RDMs and feedback history model RDMs, controlling for the influence of the capture distance RDM. **d**, Group-level partial correlation RSA results for the capture distance model, controlling for either the Recent-2 Trial or Recent-3 Trial model. *: *P* < 0.05; **: *P* < 0.01; ***: *P* < 0.001. Discovery FC Clusters ∩ Validation FC Clusters

Given the observed relationship between target object presence and feedback scores (Fig. 2a–b), and the likelihood that the appearance of these objects depends on participants’ spatial positions (e.g., traffic lights typically appear near crosswalks), we explored whether spatial location confounded the neural encoding of feedback history. Visual inspection of feedback distribution maps suggested spatial clustering of high (blue) and low (red) feedback scores across capture sites (Fig. 4b). This observation raised the possibility that the capture distance model and feedback history models may be collinear (Supplementary Table 3). Indeed, the capture distance RDM showed stronger mean RSA correlations than any of the feedback history models; for instance, the MiOG cluster in the discovery group yielded a mean *z*-scored correlation of *Z* = 0.282 for the capture distance model, compared to *Z* = 0.095 for both the Recent-2 and Recent-3 Trial feedback models.

To formally disentangle these contributions, we performed partial correlation analyses in the second-level RSA by regressing out the effects of the capture distance model or feedback history models from both neural and model RDMs. Even after controlling for capture distance, both clusters showed significant and robust encoding of the Recent-2 Trial model (MiOG: discovery, *t*(15) = 2.407, *P* = 0.017, *d* = 0.621; validation, *t*(15) = 3.354, *P* = 0.002, *d* = 0.866; IFG: discovery, *t*(15) = 2.702, *P* = 0.010, *d* = 0.698; validation, *t*(15) = 4.315, *P* < 0.001, *d* = 1.114) and the Recent-3 Trial model (IFG: discovery, *t*(15) = 3.211, *P* = 0.003, *d* = 0.829; validation, *t*(15) = 3.798, *P* = 0.001, *d* = 0.981) (Fig. 4c; Supplementary Table 4).

Conversely, when controlling for the Recent-2 and Recent-3 Trial models, the capture distance model remained significantly associated with both clusters (Fig. 4d; Supplementary Table 4). Specifically, the MiOG showed strong encoding of capture distance (discovery: *t*(15) = 7.670, *P* < 0.001, *d* = 1.980; validation: *t*(15) = 6.502, *P* < 0.001, *d* = 1.679), as did the IFG (discovery: *t*(15) = 4.850, *P* < 0.001, *d* = 1.252; validation: *t*(15) = 5.689, *P* < 0.001, *d* = 1.469), after controlling for the Recent-2 Trial model. Similar effects were observed after regressing out the Recent-3 Trial model (MiOG: discovery, *t*(15) = 7.886, *P* < 0.001, *d* = 2.036; validation, *t*(15) = 5.941, *P* < 0.001, *d* = 1.534; IFG: discovery, *t*(15) = 4.870, *P* < 0.001, *d* = 1.258; validation, *t*(15) = 5.456, *P* < 0.001, *d* = 1.409).

### The MiOG (vmPFC) and IFG (vlPFC) interact with distinct functional networks during exploration

Although the MiOG and IFG both encoded task history representations during the *Photographer* paradigm, they may participate in distinct functional networks during task performance. To examine this possibility, we conducted task-based functional connectivity (FC) analyses using the CONN toolbox, with the MiOG and IFG clusters defined from RSA as seed regions (see Methods, “*Task-based functional connectivity (FC) analysis*,” for details). Because feedback phases lasted only two seconds per trial, the analysis was restricted to fMRI volumes acquired during the exploration phases. For each subgroup, seed-based connectivity maps were generated and thresholded at the voxel level (*P* < 0.001) and corrected for multiple comparisons at the cluster level using FDR (*P* < 0.05). Conjunction FC maps between the subgroups were computed following the same approach used in the RSA pipeline.

The conjunction FC analysis revealed distinct functional networks associated with the two seed regions during exploration. The MiOG exhibited broad connectivity with the bilateral medial prefrontal cortex (mPFC), orbitofrontal cortex (OFC), anterior cingulate cortex (ACC), insula, middle temporal gyrus (MiTG), temporal pole, posterior cingulate cortex (PCC), precuneus, and angular gyrus: regions collectively comprising the default mode network (DMN; Fig. 5a). Additionally, the MiOG was functionally coupled with subcortical structures such as the VS, hippocampus, and amygdala, as well as the cerebellar Crus I/II. In contrast, the IFG showed more spatially restricted connectivity, primarily involving bilateral IFG, medial frontal gyrus, MiTG, left dorsolateral prefrontal cortex (dlPFC), and angular gyrus: components of the frontoparietal network (FPN; Fig. 5b). Connectivity was also observed in the left caudate and right-lateralized cerebellar Crus I/II.

**Fig. 5.**
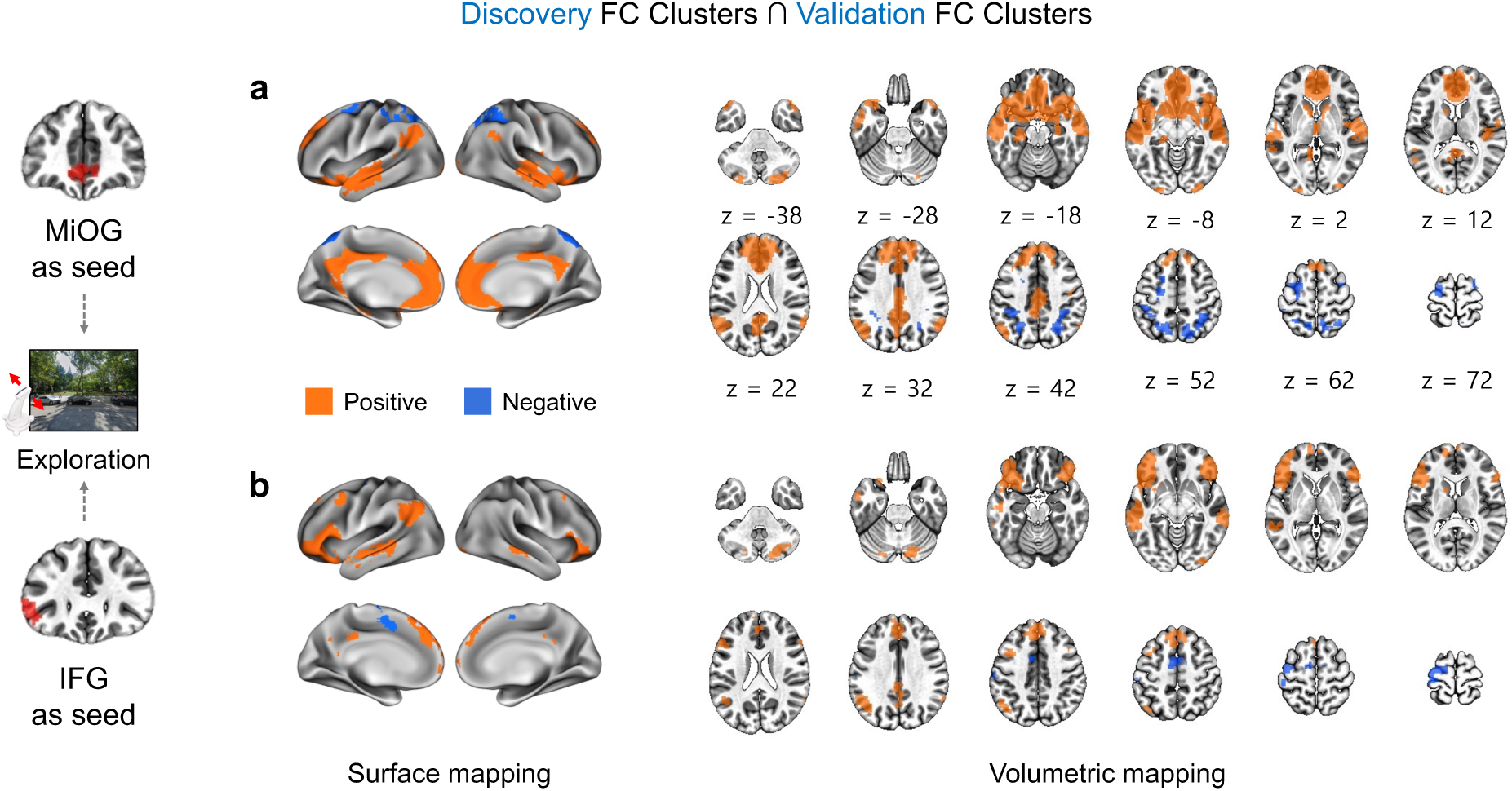
Cross-validated functional connectivity networks of MiOG and IFG during the exploration phase. Task-based functional connectivity (FC) analyses were conducted using the MiOG and IFG clusters as seed regions, focusing specifically on the exploration blocks. Following a procedure similar to the searchlight RSA pipeline, group-level FC maps were generated separately for each subgroup (voxel-level threshold *P* < 0.001; cluster-level FDR-corrected *P* < 0.05). Conjunction analyses were then applied to identify functional networks consistently coactivating with the seed regions across both groups. **a**, Surface and volumetric representations of the MiOG-seeded functional network. In both surface and volumetric plots, orange indicates positive correlations and blue indicates negative correlations. **b**, Cross-validated functional network coactivating with the IFG.

These findings suggest that the MiOG and IFG are functionally integrated with the DMN and FPN, respectively, during exploration. However, given the possibility that such couplings may reflect intrinsic connectivity rather than task-evoked processes, we performed a control analysis using data from nonexploration phases. FC maps computed during nonexploration blocks also showed connectivity patterns consistent with the DMN (for the MiOG seed) and FPN (for the IFG seed), but with markedly reduced spatial extents (Supplementary Fig. 4). Notably, the MiTG and angular gyrus clusters, which were coactivated with both seed regions during exploration, were absent in the nonexploration FC maps. Moreover, these components demonstrated significantly stronger functional coupling with the RSA-defined clusters during exploration compared to the nonexploration phase (Supplementary Figs. 5 and 6), supporting the task-specific modulation of these networks.

## Discussion

We designed a novel naturalistic RL task paradigm in which participants explored and interacted with complex, real-world street-view scenes while undergoing fMRI scanning. We identified neural correlates of this real-world learning behavior by analyzing the relationship between received feedback and neural activation at the time of feedback presentation. Interestingly, feedback scores from the current trial were not robustly represented in the brain during feedback phases. Instead, recent feedback history, which spans multiple previous trials, was robustly encoded in neural patterns within the MiOG and IFG, a finding replicated across two independent participant subgroups. These regions continued to encode feedback history and spatial location information even after controlling for potential confounding effects.

The IFG, or ventrolateral prefrontal cortex (vlPFC), is traditionally associated with speech production and sentence comprehension ^19^. Given its anatomical connections with the inferotemporal cortex ^20^, the IFG may also support working memory processes. The anterior vlPFC (Brodmann Area 47), which overlaps with our IFG cluster, has been implicated in the retrieval of semantic ^21–23^ and episodic memory ^23–25^, suggesting that it may support the retrieval of prior feedback information or internal task models.

Our second-level RSA showed that the IFG encoded spatial relationships between photographs. Although the right vlPFC is more commonly linked to spatial cognition, the bilateral vlPFC, including the left side, has been associated with route planning ^26^, replanning ^27^, and path computation under increased planning demands ^28^. Given the demands of our Photographer task, which requires continuous exploration and replanning, participants likely relied on cognitive maps stored in the vlPFC to guide decision-making.

The encoding of both feedback history and spatial maps in the IFG suggests that this region processes task-related representations in a flexible and integrative manner. Bilateral vlPFC has been linked to latent task structures that generalize across stimuli ^29^ and to model-based representations in RL environments ^30^. These findings support the idea that the vlPFC may be engaged in updating and maintaining task-specific cognitive models during feedback-based learning.

Searchlight RSA also identified a robust MiOG cluster, corresponding to the ventromedial prefrontal cortex (vmPFC), encompassing Brodmann Areas 10, 11, and 32. The vmPFC, which functionally interacts with the amygdala and ventral striatum ^31^, is well known for encoding value, particularly in the orbitofrontal cortex (OFC) ^32,33^. Despite this, our univariate analyses and trial-wise RSA did not reveal consistent value encoding in these regions, likely due to the complexity of the naturalistic environment, which impedes straightforward credit assignment ^34,35^. Participants also reported difficulty in identifying scene features associated with higher feedback scores.

Nevertheless, the vmPFC robustly encoded feedback history and spatial location information, suggesting that it may integrate feedback and contextual information across trials. For example, a participant might take a photograph near a crosswalk in Trial 3 and receive relatively high feedback. In the following trial, they focus on trees in a park, which results in a lower score. When Trial 5 again captures a crosswalk and receives high feedback, the vmPFC might associate these experiences to form a cognitive map ^33,36,37^ linking spatial contexts and feedback outcomes.

The hippocampus, traditionally linked to cognitive maps and spatial navigation ^38,39^, also showed significant associations with feedback history in our RSA, especially when using anatomically defined subregions. Second-level RSA with partial correlations revealed that all hippocampal subregions, except the left posterior hippocampus, robustly encoded both capture distance and feedback history models (Supplementary Fig. 7). These findings align with a recent hypothesis positing that the hippocampus constructs state-transition graphs, the OFC assigns values to states, and the lateral PFC stores and updates the graph ^33^. Indeed, the hippocampus showed positive connectivity with the MiOG during exploration (Supplementary Fig. 8), further supporting its role in state-value mapping.

Our FC analyses revealed distinct network affiliations for MiOG and IFG. The MiOG was predominantly connected with the DMN, including medial prefrontal cortex, posterior cingulate cortex, and temporal structures, which are implicated in internal model integration and self-guided learning ^40–43^. The DMN has also been linked to RL processes such as Q-function computation and action guidance ^44^. These findings suggest that the DMN supports exploration-based learning by integrating memory and ongoing experience.

In contrast, IFG-based connectivity maps highlighted recruitment of the FPN, including bilateral IFG and left dorsolateral prefrontal cortex (dlPFC), which are associated with executive functions such as working memory and cognitive control. These FPN components may interact with value and prediction error signals during decision-making ^45,46^. Thus, the IFG and its associated network likely contribute to the executive control of task learning.

Interestingly, the middle temporal gyrus (MiTG) and angular gyrus were coactivated with both MiOG and IFG during exploration. Given that both clusters encode feedback history and spatial information, this coactivation may reflect interactions between the DMN and FPN ^47^. These regions have also been linked to event boundary processing in naturalistic tasks ^48,49^, suggesting they may support the segmentation and integration of feedback and spatial experiences across trials.

The role of memory in task learning is also emphasized in artificial intelligence. Deep RL frameworks such as Deep Q Networks (DQNs) rely on "experience replay" buffers that store and reuse past state–action–reward–state episodes ^50^. Episodic RL models further enhance learning by using entire past episodes to compute value estimates ^51,52^. Our findings highlight the robust encoding of multi-trial feedback histories (e.g., Recent-2 and Recent-3 models), suggesting that the human brain may use a comparable strategy, leveraging stored episodic information to support learning in complex environments.

Unlike prior neuroimaging studies that employed naturalistic navigation tasks focused on wayfinding ^53,54^, our paradigm emphasized goal inference and learning from feedback with minimal instruction. This design mirrors early developmental learning, allowing us to investigate how internal models are formed through exploration. We confirmed the replicability of our neural findings across two independent subgroups and validated both RSA and FC results to ensure robustness.

One limitation of our study is the relatively small number of photographic trials, which may constrain learning opportunities. While we aimed to preserve ecological validity and limit MRI duration to one hour, future designs could balance run structure to increase trial numbers while minimizing task complexity. Additionally, although the discovery group yielded more significant RSA clusters than the validation group, further analysis revealed no systematic differences in model fit, suggesting individual variability rather than group-level effects (Supplementary Figs. 9–11).

Lastly, our study did not include formal computational models (e.g., the Rescorla‒Wagner model ^55,56^ or Bayesian RL models ^57,58^) due to the task’s complexity and limited sample size. However, deep RL models such as DQNs may be more appropriate for future research. These models can leverage convolutional or transformer-based neural networks to reduce high-dimensional input into compressed feature representations, facilitating the development of computational accounts of behavior in naturalistic environments.

In conclusion, our naturalistic RL paradigm enabled participants to learn through self-guided exploration and minimal instruction, closely simulating real-world learning. We demonstrated that both the MiOG and IFG encode multi-trial feedback history and spatial relationships, supporting cognitive mapping, memory-based learning, and executive control. These findings contribute to our understanding of human RL mechanisms and offer valuable insights for advancing AI systems inspired by human cognition.

## Methods

### Participants

The experimental protocol was approved by the Institutional Review Board at Korea University, and all participants provided written informed consent. We initially recruited 39 healthy, right-handed individuals and assigned them to one of two groups based on the year of data acquisition: the *discovery group*, whose data were collected in 2022, and the *validation group*, collected in 2023. Three participants from the validation group reported dizziness during the experiment and were excluded prior to data collection. Additionally, we excluded three participants from the discovery group and one from the validation group due to technical issues, including failures in Brain Imaging Data Structure (BIDS) conversion, errors in cortical surface reconstruction, or excessive head motion during at least one fMRI run ^59,60^. As a result, data from 32 participants were included in the final analysis, with 16 participants in each group. The discovery group included 10 females (mean age = 22.6 ± 2.2 years), and the validation group included 6 females (mean age = 23.8 ± 2.7 years) (Supplementary Table 1).

### Experimental paradigm

We developed a naturalistic, real-world RL task paradigm, termed *Photographer*, in which participants explored immersive street-view environments and engaged in a real-life task of taking photographs while undergoing fMRI scanning (Fig. 1a). The experimental environment was implemented using the Google Maps Application Programming Interface (API) (https://developers.google.com/maps), allowing us to build a dynamic street-view interface for task interaction. All task-related code was developed using React 18.2 and Electron 20.0.2.

During the experiment, participants visited five major cities—Boston, Los Angeles, New York, London, and Paris—across five task runs. Each run consisted of eight trials, with each trial comprising five sequential events: *exploration*, *capture*, *preview*, *cross-modality*, and *feedback*.

At the beginning of each trial, participants were given up to 40 seconds to freely explore the street environment using an MR-compatible joystick (Tethyx; Current Designs, Philadelphia, PA), with visual stimuli presented through MR-compatible binocular goggles. During this *exploration* phase, participants navigated the scene and were instructed to press a button to capture a scene of interest, initiating the *capture* event. If no image was taken within the time limit, the current viewpoint was automatically recorded and marked as a failed capture.

Captured images were then reviewed in a *preview* event, followed by a jittered five-second fixation. Subsequently, the *cross-modality* event presented a machine-generated description of the photograph in either text or auditory format, randomly assigned and labeled as either a *Caption* or *Voice* event. We used the Microsoft Azure AI Vision Image Analysis service (https://azure.microsoft.com/products/cognitive-services/computer-vision/) to generate captions and the Google Cloud Text-to-Speech API (https://cloud.google.com/text-to-speech) to convert text into spoken voice.

Following another jittered fixation, participants received a *feedback* event. Feedback represented the *conceptual similarity* between the captured photograph and a hidden target sentence: *“A person on the bicycle is waiting for the traffic light.”* This similarity score was computed using a multimodal deep learning model, Contrastive Language-Image Pretraining (CLIP) ^61^. CLIP uses transformer-based encoders pretrained on large-scale image–text pairs to produce embedding vectors for both images and sentences. Higher cosine similarity between the image and text embeddings reflects greater semantic alignment. We employed a JavaScript implementation of CLIP (https://github.com/josephrocca/openai-clip-js), which enabled seamless inference within the Electron environment. The raw similarity values (ranging from 0.18 to 0.28) were linearly rescaled to a 0 – 100 scale and displayed as the width of a horizontal feedback bar.

Participants were instructed to take photographs that would yield the highest possible feedback scores. However, they were not informed of the underlying feedback mechanism, the use of CLIP, or the fact that the same target sentence remained fixed across all five runs.

### Behavioral data analyses

#### Validation of the feedback score

We annotated the presence or absence of the three target objects mentioned in the target sentence (*person*, *bicycle*, and *traffic light*) in each captured image using a deep learning-based object detection model, You Only Look Once version 5 (YOLOv5) (https://github.com/ultralytics/yolov5). To evaluate the relationship between object presence and feedback, we conducted two-sample *t*-tests comparing feedback scores between images that contained versus did not contain each target object. Additionally, a one-way analysis of variance (ANOVA) was performed to examine the main effect of the number of target objects present in a scene on the corresponding feedback score.

All statistical analyses were conducted in R version 4.2.2, using the *rstatix* package for hypothesis testing ^62^ and the *effectsize* package for estimating effect sizes ^63^.

#### Validation of learning

To assess systematic changes in feedback scores across task runs, we employed linear mixed-effects (LME) models separately for the discovery and validation groups. Model fitting was conducted using the *lmerTest* package ^64^ in R version 4.2.2. The between-run effect model included the fixed effect of task Run, along with random intercepts for both trial and subject:

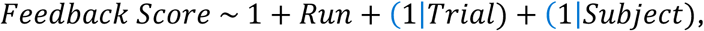

where *Run* denotes the run number, and (1|Trial) and (1|Subject) specify the random intercepts for trial and subject, respectively. Model fit was evaluated by comparing log-likelihood improvements over a null model (random intercepts only) using one-way ANOVA and differences in Akaike Information Criterion (AIC). Post hoc *t*-tests comparing estimated marginal means across runs were performed using the *emmeans* package ^65^, with resulting *p*-values corrected for multiple comparisons using the false discovery rate (FDR) method.

We hypothesized that participants might gradually learn the hidden task structure over the five runs, despite exploring different environments across five cities. Given the high trial-to-trial variability in feedback scores, we computed moving-averaged scores using a sliding window of eight trials. To model group-level learning trajectories across these capture attempts, we fitted five polynomial LME models using the global trial index (GTI), defined as the chronological order of trials from the 8^th^ to the 40^th^ trial, as a predictor of the moving average feedback score (MA₈):

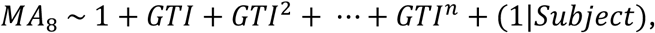

where *MA₈* is the moving-averaged feedback score over an eight-trial window, and *n* ranges from 0 (intercept-only model) to 4 (quartic model). The best-fitting model for each group was selected based on improvements in log-likelihood and AIC.

#### MRI data acquisition

Structural and functional MRI data were acquired using a 3.0 T Siemens Tim-Trio scanner equipped with a 12-channel head coil. High-resolution T1-weighted structural images were obtained using a magnetization-prepared rapid gradient echo (MPRAGE) sequence with the following parameters: repetition time (TR) = 1,900 ms; echo time (TE) = 2.52 ms; flip angle (FA) = 9°; 176 sagittal slices with no interslice gap; field of view (FoV) = 256 × 256 mm²; and isotropic voxel size of 1 × 1 × 1 mm³.

*T2*\**-weighted functional images* were acquired using a standard gradient-echo echo-planar imaging (EPI) sequence, with the following parameters: TR = 2,000 ms; TE = 30 ms; FA = 84°; 35 contiguous axial slices; FoV = 240 × 240 mm²; and voxel size = 3 × 3 × 4 mm³.

#### MRI data preprocessing

All structural and functional MR images were converted into the BIDS format using HeuDiConv ^66^. Preprocessing of the BIDS-formatted data was conducted separately for each group using fMRIPrep version 23.0.2 ^67,68^ (RRID:SCR_016216), which is built upon Nipype version 1.8.6 ^69,70^ (RRID:SCR_002502). Detailed preprocessing steps are described in the Supplementary Methods under the section “**MRI data preprocessing via fMRIPrep**.”

#### General linear models (GLMs)

We constructed two general linear models (GLMs) for univariate (GLM 1) and multivariate (GLM 2) analyses using AFNI through the Nipype interface. Preprocessed fMRI volumes were resampled to an isotropic 3 mm voxel space. For univariate analysis (GLM 1), the images were spatially smoothed using an 8 mm FWHM Gaussian kernel and scaled to a mean BOLD signal of 100. For multivariate analysis (GLM 2), no spatial smoothing was applied.

GLM 1 estimated beta values for each task event block per run (i.e., *Exploration*, *Capture*/*Capture-failed*, *Preview*, *Caption*/*Voice*, and *Feedback*). Each task regressor included multiple onsets corresponding to specific task events across the eight trials in each run. For the *Feedback* event, an additional parametric modulator was included to model trial-by-trial feedback score fluctuations through amplitude modulation. GLM 1 also incorporated 26 nuisance regressors, including zero- to fifth-order polynomial trends for detrending, six head motion parameters and their first-order derivatives, three global signals (whole brain, cerebrospinal fluid [CSF], and white matter) and their derivatives, framewise displacement (FD), and outlier volume indices identified by fMRIPrep.

Group-level *t*-tests were performed independently for the discovery and validation groups to identify clusters encoding feedback events or feedback scores, with voxel-level thresholds of *P* < 0.005 (one-sided) and cluster-level *α* < 0.05.

GLM 2 estimated trialwise beta values for each task event. Each regressor modeled a single time point corresponding to one of five task events across the eight trials, resulting in 40 regressors (5 task events × 8 trials). Parametric modulators were not included in GLM 2, and the same set of nuisance regressors from GLM 1 was applied.

#### fMRI univariate analysis

The estimated beta maps from GLM 1 for each run were averaged across runs for each event within each participant. Group-level inference was performed using one-sample *t*-tests on these averaged beta maps across participants, implemented via AFNI’s 3dttest++ with cluster-level multiple comparison correction enabled using the -ClustSim option. The resulting statistical maps were constrained within a gray matter (GM) mask, defined by thresholding the MNI152NLin2009cAsym GM probability map at 0.3.

Significant positive clusters were identified using a voxel-level threshold of *P* < 0.005 (one-sided) and a cluster-level alpha of 0.05. We conducted group-level inference and cluster correction separately for the discovery and validation subgroups. The two resulting statistical maps were then overlapped to identify cross-validated clusters, defined as contiguous voxel regions with at least 20 voxels in both subgroups.

#### Representational similarity analysis (RSA) with fMRI

We hypothesized that feedback information is encoded in multivoxel activation patterns rather than in univariate BOLD signal fluctuations at the single-voxel level. Additionally, we examined the possibility that participants retrospectively referenced previous trials by recalling captured photographs and their associated feedback scores, to update their internal task models. To test these hypotheses, we conducted RSA by comparing multivoxel neural activation patterns with behavioral data.

### Model representational dissimilarity matrices (RDMs)

Each trial and its corresponding captured image were associated with a feedback score. To examine how these feedback scores were represented across trials, we constructed representational dissimilarity matrices (RDMs) for both single-trial and multiple-trial history models. For the single-trial model (denoted as ‘Current Trial’; Fig. 6a), we computed Euclidean distances between feedback scores of all trial pairs within a run. In contrast, the multiple-trial history models represented each trial as a vector of concatenated feedback scores from the current trial and one or two preceding trials, which forms the ‘Recent-2 Trial’ and ‘Recent-3 Trial’ models, respectively (Fig. 6b). Euclidean distances were then computed between these feedback history vectors to define the dissimilarity structure across trials.

**Fig. 6.**
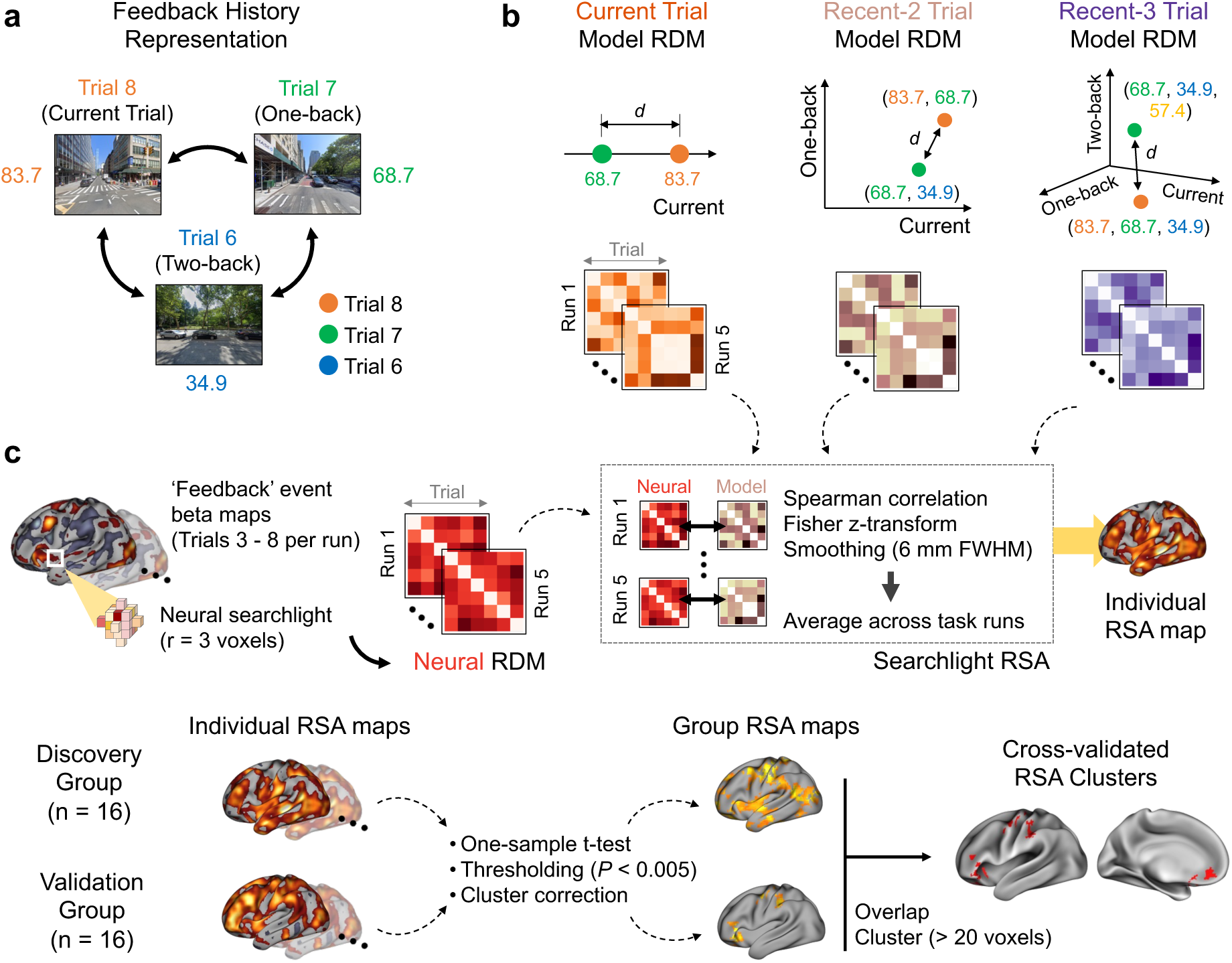
Searchlight RSA pipeline using feedback history models. **a**, Illustration of feedback history representation. During the *Photographer* task, a participant may internally track feedback from the current, one-back, and two-back trials. Comparing feedback scores across trials may facilitate inference of the task’s hidden structure. **b**, Geometric representation of three feedback history model RDMs. The Current Trial model computes Euclidean distances between trials based on their individual feedback scores in a one-dimensional (1D) axis. The Recent-2 and Recent-3 Trial models extend this by calculating distances in 2D (current + one-back) or 3D (current + one-back + two-back) spaces, respectively. To ensure fair comparison, trials 3–8 from each run were used to construct five 6×6 RDMs for each model. **c**, Searchlight RSA with cross-validation. For each participant, neural RDMs derived from feedback-related activation patterns are compared to a model RDM, and RSA results are averaged across task runs (top row). Thresholded and cluster-corrected RSA maps are generated at the group level. Overlapping significant clusters from both subgroups define cross-validated RSA clusters representing the target model RDM (bottom row).

For the single-trial history model RDMs, we defined three distinct models: the current trial, one-back trial, and two-back trial models. In these, each trial t was represented solely by the feedback score from trial t, t-1, or t-2, respectively. Euclidean distances were computed between each pair of trials in a run, using their respective feedback scores as single-point representations. To ensure comparability across models, only Trials 3 to 8 were included in the analysis, resulting in 6 × 6 RDMs for each model per run.

In the multiple-trial history RDMs, each trial t was represented by a vector of concatenated feedback scores: [t, t-1] for the Recent-2 Trial model and [t, t-1, t-2] for the Recent-3 Trial model. We computed Euclidean distances between these multidimensional vectors across all eligible trial pairs, resulting again in 6 × 6 model RDMs per run for each participant.

All multiple-trial models captured information from previous trial attempts. We hypothesized that the brain may represent aggregated feedback history more robustly than single-trial feedback. To test this, we defined three control RDMs focused solely on past feedback: the one-back and two-back single-trial models, and a “Previous-2 Trial” model that combined feedback scores from trials t-1 and t-2, excluding the current trial.

Because participants were not informed that all task runs shared the same target sentence, they may have assumed that different city contexts implied different feedback rules. This likely resulted in non-uniform feedback distributions across runs. Accordingly, all model RDMs were computed separately for each task run and participant to account for potential run-specific variations.

### Neural searchlight RSA

We employed trialwise beta maps of feedback events from GLM 2, where internal model updates are most likely to occur during feedback score presentation. To improve the reliability of neural RDMs, we applied univariate noise normalization by dividing voxelwise beta values by the standard deviations of the GLM 2 residuals ^71^. Neural RDMs were constructed using a searchlight approach with three-voxel radius spheres traversing the gray matter (GM) mask. For each sphere, we computed correlation distances (1 – Pearson’s *r*) between beta patterns across trials, resulting in a 6 × 6 neural RDM for each task run.

For each model RDM (representing different feedback history structures), we vectorized the upper triangular elements of both the model and neural RDMs and calculated the rank-based correlation (Spearman’s ρₐ) between the two vectors. The resulting individual RSA maps were Fisher’s r-to-z transformed and spatially smoothed using a 6 mm FWHM Gaussian kernel. Searchlight RSA was performed separately for each task run and for each feedback model. Runwise RSA maps were averaged across runs within each participant (Fig. 6c).

We conducted group-level analysis using one-sample *t* tests on the average RSA maps across participants to identify voxels that robustly encoded each feedback model across task runs. This was performed using AFNI’s 3dttest++ with cluster-level multiple comparison correction (voxel-level P < 0.005, one-sided; cluster-level α < 0.05). Independent searchlight RSA and group-level tests were repeated for the discovery and validation groups. We then overlapped the two thresholded statistical maps and identified cross-validated clusters (≥ 20 voxels) that consistently represented a given feedback model across both groups.

#### Second-level RSA

We investigated whether additional task-related information was encoded within the clusters that robustly represented feedback history models. In addition to the six feedback history model RDMs derived from previous RSAs (i.e., Current Trial, One-back Trial, Two-back Trial, Recent-2 Trial,

Recent-3 Trial, and Previous-2 Trial), we constructed three exploration-related model RDMs: Exploration Time, Exploration Distance, and Capture Distance. The Exploration Time model captured the elapsed time between the trial onset and the capture action, comparing exploration durations across trials. The Exploration Distance model represented geodesic distances (computed using the Haversine formula; https://www.movable-type.co.uk/scripts/latlong.html) between the exploration starting point and the final capture location for each trial. The Capture Distance model compared trialwise geodesic distances between capture locations, forming a spatial map of all captured photographs within each run.

All model RDMs were compared to neural RDMs computed from trialwise feedback beta values within each previously identified cluster mask. We followed the same RSA procedure described in the “*Neural Searchlight RSA*” section, except that no spatial smoothing was applied. For group-level inference, we conducted permutation-based one-sample *t* tests (10,000 permutations) for each model RDM to determine statistical significance.

Given the observed potential confounding relationship between the Capture Distance model and the feedback history models, we performed partial RSA to control for these effects. Specifically, we regressed out the influence of the confounding model from both the neural and model RDMs and computed RSA on the resulting residuals. The resulting partial correlations were Fisher r-to-z transformed and subjected to one-sample *t* tests using 10,000 permutations.

In summary, we examined four main neural-model relationships: (1) Neural RDM vs. Recent-2 Trial RDM, controlling for the Capture Distance RDM; (2) Neural RDM vs. Recent-3 Trial RDM, controlling for the Capture Distance RDM; (3) Neural RDM vs. Capture Distance RDM, controlling for the Recent-2 Trial RDM; and (4) Neural RDM vs. Capture Distance RDM, controlling for the Recent-3 Trial RDM.

#### Task-based functional connectivity (FC) analysis

Functional connectivity (FC) preprocessing and denoising were performed using the CONN toolbox ^72^. For details, please refer to the Supplementary Methods section, “**Functional Connectivity (FC) Analysis via CONN**.”

### First-level FC analysis

Seed-based FC maps were estimated to characterize the connectivity patterns associated with the MiOG and IFG seed regions identified via searchlight RSA. FC strength was quantified using Fisher-transformed bivariate correlation coefficients derived from a weighted general linear model (weighted GLM) ^73^. This model was computed separately for each seed–target voxel pair, capturing the temporal correlation between their respective BOLD time series. Individual scans were weighted using a boxcar regressor corresponding to each task or experimental condition, convolved with the SPM canonical hemodynamic response function and subsequently rectified.

### Group-level FC analysis

After estimating voxel-level FC values for each participant, group-level inference was conducted using multivariate parametric statistics, incorporating random effects across subjects and sample covariance estimation across multiple measurements. Inferences were made at the cluster level, based on groups of contiguous voxels, using parametric statistics derived from Gaussian random field theory ^73,74^. Multiple comparison correction was applied to generate group-level inference maps using a cluster-forming voxel-level threshold of *P* < 0.001 and a familywise error-corrected cluster-size threshold of *P*FDR < 0.05 ^75^.

To identify connectivity patterns that were robust across subgroups, we performed conjunction analyses on the group inference maps from the discovery and validation groups. Specifically, we first separated positively and negatively correlated clusters from each group’s map, intersected the respective positive or negative masks between groups, and then merged them to construct cross-validated FC maps for each seed region.

Since the observed FC patterns may not be specific to the exploration phase, we repeated the first- and group-level analyses using fMRI volumes corresponding to nonexploration periods (i.e., cross-fixation, preview, cross-modal, and feedback blocks) with the same seed regions. This control analysis revealed additional cross-validated functional networks associated with the nonexploration phase. To assess phase-specific connectivity, we also conducted paired *t*-tests between exploration and nonexploration FC maps for each subgroup (voxel-level threshold: *P* < 0.001; cluster-level threshold: *P*FDR < 0.05).

## Reporting summary

Further information on the research design is available in the Nature Portfolio Reporting Summary linked to this article.

## Data availability

All neuroimaging and behavioral datasets employed in this study will be available upon reasonable request to the corresponding author.

## Code availability

The Photographer Paradigm is available at https://github.com/constantjin/photographer-experimental-paradigm. Analysis scripts for first-/second-level RSA and visualization are open to the public at https://github.com/constantjin/photographer-data-analysis.

## Acknowledgments

The authors thank Mr. Minseok Choi for his logistic support during the data collection and helpful discussion. This work was supported by a National Research Foundation (NRF) grant funded by the Korean government (MSIT) (NRF-2021M3E5D2A01022515, No. RS-2023-00218987; RS-2025-00562405) and, in part, by an Electronics and Telecommunications Research Institute (ETRI) grant funded by the Korean government [25ZS1100, Core Technology Research for Self-Improving Integrated Artificial Intelligence System]. The funders had no role in the study design, data collection, analysis or interpretation of the data, manuscript preparation, or decision to submit for publication.

## Author contributions

Conceptualization: S.J., J.L. and J.H.L. Data Collection: S.J. and J.L. Methodology: S.J., J.L. and J.H.L. Software: S.J. Formal Analysis: S.J. Resources: J.H.L. Data Curation: S.J. Writing - Original Draft: S.J. and J.H.L. Writing - Review & Editing: S.J., J.L. and J.H.L. Visualization: S.J. Supervision: J.H.L. All the authors approved the final version of the manuscript.

