## Supplementary Materials and Methods for "A naturalistic reinforcement learning task uncovers historical neural representations"

### Supplementary Methods

#### MRI data preprocessing via fMRIPrep

The following preprocessing description is adapted from the automatically generated boilerplate text provided by fMRIPrep.

*Anatomical MRI data preprocessing*
A total of one $T_1$-weighted (T1w) image was found within the input BIDS dataset. The T1w image was corrected for intensity nonuniformity (INU) with N4BiasFieldCorrection[1], distributed with ANTs 2.3.3[2] (RRID:SCR_004757), and used as a T1w reference throughout the workflow. The T1w-reference was then skull-stripped with a Nipype implementation of the antsBrainExtraction.sh workflow (from ANTs), using OASIS30ANTs as the target template. Brain tissue segmentation of cerebrospinal fluid (CSF), white matter (WM) and gray matter (GM) was performed on the brain-extracted T1W image via fast[3] (FSL 6.0.5.1:57b01774, RRID:SCR_002823). The brain surfaces were reconstructed via recon-all[4] (FreeSurfer 7.3.2, RRID:SCR_001847), and the brain mask estimated previously was refined with a custom variation of the method to reconcile ANT-derived and FreeSurfer-derived segmentations of the cortical gray matter of Mindboggle[5] (RRID:SCR_002438). Volume-based spatial normalization to one standard space (MNI152NLin2009cAsym) was performed through nonlinear registration with antRegistration (ANTs 2.3.3) via brain-extracted versions of both the T1w reference and the T1w template. The following template was selected for spatial normalization and accessed with TemplateFlow[6] (23.0.0): ICBM 152 Nonlinear Asymmetrical template version 2009c[7] [RRID:SCR_008796; TemplateFlow ID: MNI152NLin2009cAsym].

*Functional MRI data preprocessing*
For each of the 5 BOLD fMRI runs found per subject (across all tasks and sessions), the following preprocessing was performed. First, a reference volume and its skull-stripped version were generated via the custom methodology of fMRIPrep. Head motion parameters with respect to the BOLD reference (transformation matrices and six corresponding rotation and translation parameters) are estimated before any spatiotemporal filtering using mcflirt[8] (FSL 6.0.5.1:57b01774). BOLD runs were slice-time corrected to 0.975 s (0.5 of slice acquisition range 0–1.95 s) via 3dTshift from AFNI[9] (RRID:SCR_005927). The BOLD time series (including slice-timing correction when applied) were resampled onto their original, native space by applying the transforms to correct for head motion. These resampled BOLD time series will be referred to as preprocessed BOLD signals in the original space or just preprocessed BOLD signals. The BOLD reference was then coregistered to the T1w reference via bbregister (FreeSurfer), which implements boundary-based registration[10]. Coregistration was configured with six degrees of freedom. Several confounding time series were calculated on the basis of the preprocessed BOLD signal: framewise displacement (FD), DVARS and three regionwise global signals. FD was computed via two formulations: power[11] (absolute sum of relative motions) and Jenkinson[8] (relative root mean square displacement between affines). FD and DVARS are calculated for each functional run, both of which use their implementations in Nipype (following the definitions by Power *et al.* 2014[11]). The three global signals are extracted within the CSF, the WM, and the whole-brain masks. Additionally, a set of physiological regressors was extracted to allow for component-based noise correction[12] (CompCor). Principal components are estimated after high-pass filtering of the preprocessed BOLD time series (using a discrete cosine filter with a 128 s cutoff) for the two CompCor variants: temporal (tCompCor) and anatomical (aCompCor). tCompCor components are then calculated

from the top 2% of the variable voxels within the brain mask. For aCompCor, three probabilistic masks (CSF, WM, and combined CSF+WM) are generated in anatomical space. The implementation differs from that of Behzadi *et al.* in that instead of eroding the masks by 2 pixels on BOLD space, a mask of pixels that likely contain a volume fraction of GM is subtracted from the aCompCor masks. This mask is obtained by dilating a GM mask extracted from FreeSurfer's average segmentation, and it ensures that the components are not extracted from voxels containing a minimal fraction of the GM. Finally, these masks are resampled into BOLD space and binarized by thresholding at 0.99 (as in the original implementation). The components are also calculated separately within the WM and CSF masks. For each CompCor decomposition, the k components with the largest singular values are retained such that the retained components' time series are sufficient to explain 50% of the variance across the nuisance mask (CSF, WM, combined, or temporal). The remaining components are excluded from consideration.

The head motion estimates calculated in the correction step were also placed within the corresponding confounds file. The confounding time series derived from head motion estimates and global signals were expanded with the inclusion of temporal derivatives and quadratic terms for each[13]. Frames that exceeded a threshold of 0.5 mm FD or 1.5 standardized DVARS were annotated as motion outliers. Additional nuisance time series are calculated via principal component analysis of the signal found within a thin band (crown) of voxels around the edge of the brain, as proposed by (Patriat, Reynolds, and Birn 2017)[14]. The BOLD time series were resampled into standard space, generating a preprocessed BOLD run in MNI152NLin2009cAsym space. First, a reference volume and its skull-stripped version were generated via the custom methodology of fMRIPrep. All resamplings can be performed with a single interpolation step by composing all the pertinent transformations (i.e., head–motion transform matrices, susceptibility distortion correction when available, and coregistrations to anatomical and output spaces). Gridded (volumetric) resamplings were performed via antApply Transforms (ANTs), which were configured with Lanczos interpolation to minimize the smoothing effects of other kernels[15]. Nongridded (surface) resamplings were performed via mri_vol2surf (FreeSurfer). Many internal operations of fMRIPrep use Nilearn 0.9.1 (Abraham *et al.* 2014, RRID:SCR_001362), mostly within the functional processing workflow. For more details of the pipeline, see the section corresponding to workflows in fMRIPrep's documentation (https://fmriprep.readthedocs.io/en/latest/workflows.html).

**Functional connectivity (FC) analysis via CONN**

The following description is adapted from the boilerplate text provided by the CONN [16] (RRID:SCR_009550) release 22.a [17] using SPM [18] (RRID:SCR_007037) release 12.7771, and has been updated to reflect the specifics of our data analyses.

*Preprocessing*

Functional and anatomical data were preprocessed via a modular preprocessing pipeline[19,] including realignment with correction of susceptibility distortion interactions, slice timing correction, outlier detection, direct segmentation and MNI space normalization, and smoothing. The functional data were realigned via SPM (realign & unwarp procedure)[20], where all the scans were coregistered to a reference image (first scan of the first session) via a least squares approach and a six-parameter (rigid body) transformation[21] and resampled via b-spline interpolation to correct for motion and magnetic susceptibility interactions. Temporal misalignment between different slices of the functional data was corrected following the SPM slice-timing correction procedure[22,23], which uses sinc temporal interpolation to resample each slice BOLD time series to a common mid-acquisition time. Potential outlier scans were identified via the ART toolbox[24] for acquisitions with framewise displacement above 0.9 mm or global BOLD signal changes above 5 standard deviations[11,25], and a reference BOLD image was computed for each subject by averaging all scans excluding outliers. The functional and anatomical data were normalized into standard MNI space, segmented into gray matter, white matter, and CSF tissue classes, and resampled to 3 mm isotropic voxels following a direct normalization procedure[25,26] via the SPM unified segmentation and normalization algorithm[27,28] with the default IXI-549 tissue probability map template. Finally, the functional data were smoothed via spatial convolution with an 8 mm FWHM Gaussian kernel.

*Denoising*

fMRI data were denoised via a standard denoising pipeline[19,] including the regression of potential confounding effects characterized by white matter time series (5 CompCor noise components), CSF time series (5 CompCor noise components), motion parameters and their first-order derivatives (12 factors)[29], outlier scans (below 65 factors)[11], session effects and their first-order derivatives (2 factors), and linear trends (2 factors) within each functional run, followed by bandpass frequency filtering of the BOLD time series[30] between 0.008 Hz and 0.09 Hz. The CompCor[12,31] noise components within white matter and CSF were estimated by computing the average BOLD signal as well as the largest principal components orthogonal to the BOLD average, motion parameters, and outlier scans within each subject's eroded segmentation masks. From the number of noise terms included in this denoising strategy, the effective degrees of freedom of the BOLD signal after denoising were estimated to range from 211.9–349 (average of 293.5) across the discovery group and from 205.7–326.4 (average of 286.4) across the validation group[25].

**Supplementary Tables**

**Supplementary Table 1. Participant characteristics in the discovery and validation groups**.

| | Year | N | Gender | Age (mean±s.d.) | MDBF (mean±s.d.) | VAS (mean±s.d.) | EHI |
|---|---|---|---|---|---|---|---|
| Discovery | 2022 | 16 | 6 M/ 10 F | 22.6 ± 2.2 | 59.9 ± 7.7 | 3.8 ± 2.7 | 100% |
| Validation | 2023 | 16 | 10 M/ 6 F | 23.8 ± 2.7 | 58.1 ± 7.4 | 4.5 ± 2.0 | 100% |
| Subgroup Difference | | | | $t(28.9) = -1.29$ ($P > 0.1$) | $t(30.0) = 0.65$ ($P > 0.1$) | $t(28.1) = -0.85$ ($P > 0.1$) | |

N, number of participants; MDBF, Multidimensional Mood State Questionnaire[32]; BAI, Beck Anxiety Inventory[33]; VAS, Visual Analog Scale for stress perception[34]; EHI, Edinburgh Handedness Inventory[35].

**Supplementary Table 2. Cross-validated searchlight RSA clusters for multiple-trial feedback history models.** Individual RSA maps for each subgroup were tested using a one-sample t-test with cluster-level correction (voxel-level P < 0.005, one-sided; cluster-level α < 0.05). Group-level statistical maps were then overlapped and thresholded to include only clusters with a minimum size of 20 voxels. Anatomical labels were identified using the Eickhoff–Zilles macro labels from the N27 (CA_N27_ML) atlas [36] via AFNI's 'whereami' program, reporting only those with more than 5% overlap.

| CA_N27_ML Atlas | Cluster extent (voxels) | Center-of-Mass MNI coordinates (mm) | | |
|---|---|---|---|---|
| | | x | y | z |
| ***Recent-2 Trial model*** | | | | |
| Rectal Gyrus (L/R), Mid Orbital Gyrus (L/R), Superior Orbital Gyrus (L) | 168 | –2 | 42 | –8 |
| Inferior Frontal Gyrus pars Triangularis (L), Inferior Frontal Gyrus pars Orbitalis (L) | 154 | –47 | 32 | 2 |
| ***Recent-3 Trial model*** | | | | |
| Inferior Frontal Gyrus pars Orbitalis (L), Olfactory Cortex (L), Rectal Gyrus (L) | 455 | –28 | 22 | –7 |
| Precentral Gyrus (L), Inferior Parietal Lobule (L), Postcentral Gyrus (L), Supramarginal Gyrus (L) | 224 | –47 | –19 | 49 |
| Mid Orbital Gyrus (L/R), Rectal Gyrus (L/R), Superior Orbital Gyrus (L) | 220 | 2 | 42 | –7 |
| Caudate Nucleus (R), Putamen (R), Rectal Gyrus (R) | 154 | 17 | 15 | 7 |
| Precentral Gyrus (L), Postcentral Gyrus (L) | 35 | –31 | -26 | 60 |
| ***Recent-2 Trial ∩ Recent-3 Trial*** | | | | |
| Rectal Gyrus (L/R), Mid Orbital Gyrus (L/R), Superior Orbital Gyrus (L) | 126 | –1 | 42 | –7 |
| Inferior Frontal Gyrus pars Orbitalis (L). Inferior Frontal Gyrus pars Triangularis (L) | 97 | –50 | 31 | –1 |

**Supplementary Table 3. Group-level analysis of associations between the capture distance model and feedback history models**. To assess potential confounding between the capture distance model RDM and feedback history model RDMs, we performed RSA by computing z-scored Spearman correlations between the two model types for each participant. One-sample *t*-tests with 10,000 random permutations were then applied to test each feedback history model. Additionally, two-sample *t*-tests with 10,000 permutations were used to compare RSA correlations between the two subgroups, testing for systematic group-level differences. Significant confounding effects with the capture distance model were observed for the Current Trial and Recent-3 Trial models ($P < 0.05$, one-sided). However, no significant subgroup-level differences were found across models ($P > 0.1$, two-sided), though the Recent-3 Trial and Previous-2 Trial models showed relatively larger effect sizes. μ: mean RSA correlation across participants; *: $P < 0.05$; **: $P < 0.01$; ***: $P < 0.001$.

| Feedback History RDM | Subgroup | | Subgroup Difference |
|---|---|---|---|
| | Discovery | Validation | |
| Current Trial | $\mu$ = 0.093, $t(15)$ = 2.740<br>$P$ = 0.009**, $d$ = 0.707 | $\mu$ = 0.076, $t(15)$ = 1.917<br>$P$ = 0.037*, $d$ = 0.495 | $P > 0.1$, $d$ = 0.113 |
| One-back Trial | $\mu$ = 0.030, $t(15)$ = 0.862<br>$P > 0.1$, $d$ = 0.223 | $\mu$ = -0.009, $t(15)$ = -0.317<br>$P > 0.1$, $d$ = -0.082 | $P > 0.1$, $d$ = 0.304 |
| Two-back Trial | $\mu$ = 0.037, $t(15)$ = 1.069<br>$P > 0.1$, $d$ = 0.276 | $\mu$ = 0.003, $t(15)$ = 0.091<br>$P > 0.1$, $d$ = 0.023 | $P > 0.1$, $d$ = 0.247 |
| Recent-2 Trial | $\mu$ = 0.110, $t(15)$ = 3.933<br>$P < 0.001$***, $d$ = 1.016 | $\mu$ = 0.062, $t(15)$ = 1.665<br>$P$ = 0.058, $d$ = 0.430 | $P > 0.1$, $d$ = 0.369 |
| Recent-3 Trial | $\mu$ = 0.162, $t(15)$ = 4.302<br>$P < 0.001$***, $d$ = 1.111 | $\mu$ = 0.078, $t(15)$ = 2.141<br>$P$ = 0.026*, $d$ = 0.553 | $P > 0.1$, $d$ = 0.571 |
| Previous-2 Trial | $\mu$ = 0.097, $t(15)$ = 2.328<br>$P$ = 0.017*, $d$ = 0.601 | $\mu$ = 0.015, $t(15)$ = 0.505<br>$P > 0.1$, $d$ = 0.130 | $P > 0.1$, $d$ = 0.575 |

**Supplementary Table 4. Partial correlation RSA for prefrontal and hippocampal clusters.** Neural RDMs were constructed from trial-wise beta values at feedback events for each prefrontal cluster identified by the first-level searchlight RSA (MiOG and IFG) and for hippocampal regions segmented by hemisphere (left/right) and axis (anterior/posterior)[37]. These neural RDMs were compared to model RDMs (listed under 'Model RDM') after regressing out the influence of control RDMs ('Control RDM') from both the neural and model RDMs. Individual RSA correlations were Fisher z-transformed and tested using one-sample permutation *t*-tests (10,000 permutations, one-sided). All clusters significantly encoded the capture distance model in both subgroups. In addition, all clusters, except the left posterior hippocampus, robustly represented one or more feedback history models. μ: mean RSA correlation across participants; *: $P < 0.05$; **: $P < 0.01$; ***: $P < 0.001$.

| Model RDM | Control RDM | Subgroup: Discovery | Subgroup: Validation |
|---|---|---|---|
| ***MiOG*** | | | |
| Recent-2 Trial | Capture Distance | $\mu$ = 0.067, $t(15)$ = 2.407<br>$P$ = 0.017*, $d$ = 0.621 | $\mu$ = 0.115, $t(15)$ = 3.354<br>$P$ = 0.002**, $d$ = 0.866 |
| Recent-3 Trial | Capture Distance | $\mu$ = 0.048, $t(15)$ = 1.263<br>$P$ > 0.1, $d$ = 0.326 | $\mu$ = 0.107, $t(15)$ = 3.111<br>$P$ = 0.004**, $d$ = 0.803 |
| Capture Distance | Recent-2 Trial | $\mu$ = 0.267, $t(15)$ = 7.670<br>$P$ < 0.001***, $d$ = 1.980 | $\mu$ = 0.285, $t(15)$ = 6.502<br>$P$ < 0.001***, $d$ = 1.679 |
| Capture Distance | Recent-3 Trial | $\mu$ = 0.257, $t(15)$ = 7.886<br>$P$ < 0.001***, $d$ = 2.036 | $\mu$ = 0.269, $t(15)$ = 5.941<br>$P$ < 0.001***, $d$ = 1.534 |
| ***IFG*** | | | |
| Recent-2 Trial | Capture Distance | $\mu$ = 0.064, $t(15)$ = 2.702<br>$P$ = 0.010**, $d$ = 0.698 | $\mu$ = 0.115, $t(15)$ = 4.315<br>$P$ < 0.001***, $d$ = 1.114 |
| Recent-3 Trial | Capture Distance | $\mu$ = 0.073, $t(15)$ = 3.211<br>$P$ = 0.003**, $d$ = 0.829 | $\mu$ = 0.131, $t(15)$ = 3.798<br>$P$ = 0.001**, $d$ = 0.981 |
| Capture Distance | Recent-2 Trial | $\mu$ = 0.185, $t(15)$ = 4.850<br>$P$ < 0.001***, $d$ = 1.252 | $\mu$ = 0.227, $t(15)$ = 5.456<br>$P$ < 0.001***, $d$ = 1.469 |
| Capture Distance | Recent-3 Trial | $\mu$ = 0.172, $t(15)$ = 4.870<br>$P$ < 0.001***, $d$ = 1.258 | $\mu$ = 0.229, $t(15)$ = 5.456<br>$P$ < 0.001***, $d$ = 1.409 |
| ***Hippocampus (Anterior, Left)*** | | | |
| Recent-2 Trial | Capture Distance | $\mu$ = 0.080, $t(15)$ = 2.317<br>$P$ = 0.015*, $d$ = 0.598 | $\mu$ = 0.089, $t(15)$ = 2.577<br>$P$ = 0.013*, $d$ = 0.665 |
| Recent-3 Trial | Capture Distance | $\mu$ = 0.060, $t(15)$ = 1.608<br>$P$ = 0.068, $d$ = 0.415 | $\mu$ = 0.095, $t(15)$ = 4.210<br>$P$ < 0.001***, $d$ = 1.087 |
| Capture Distance | Recent-2 Trial | $\mu$ = 0.265, $t(15)$ = 5.779<br>$P$ < 0.001***, $d$ = 1.492 | $\mu$ = 0.235, $t(15)$ = 5.502<br>$P$ < 0.001***, $d$ = 1.421 |
| Capture Distance | Recent-3 Trial | $\mu$ = 0.259, $t(15)$ = 6.557<br>$P$ < 0.001***, $d$ = 1.693 | $\mu$ = 0.220, $t(15)$ = 5.223<br>$P$ < 0.001***, $d$ = 1.349 |
| ***Hippocampus (Anterior, Right)*** | | | |
| Recent-2 Trial | Capture Distance | $\mu$ = 0.087, $t(15)$ = 2.903<br>$P$ = 0.006**, $d$ = 0.750 | $\mu$ = 0.046, $t(15)$ = 0.995<br>$P$ > 0.1, $d$ = 0.257 |
| Recent-3 Trial | Capture Distance | $\mu$ = 0.073, $t(15)$ = 2.983<br>$P$ = 0.006**, $d$ = 0.770 | $\mu$ = 0.078, $t(15)$ = 1.793<br>$P$ = 0.045*, $d$ = 0.463 |
| Capture Distance | Recent-2 Trial | $\mu$ = 0.243, $t(15)$ = 6.800<br>$P$ < 0.001***, $d$ = 1.756 | $\mu$ = 0.268, $t(15)$ = 5.181<br>$P$ < 0.001***, $d$ = 1.338 |
| Capture Distance | Recent-3 Trial | $\mu$ = 0.219, $t(15)$ = 6.916 | $\mu$ = 0.250, $t(15)$ = 4.699 |

| | | | |
|---|---|---|---|
| | | $P < 0.001$***, $d = 1.786$ | $P < 0.001$***, $d = 1.213$ |
| ***Hippocampus (Posterior, Left)*** | | | |
| Recent-2 Trial | Capture Distance | $\mu = 0.025$, $t(15) = 0.626$<br>$P > 0.1$, $d = 0.162$ | $\mu = 0.036$, $t(15) = 0.748$<br>$P > 0.1$, $d = 0.193$ |
| Recent-3 Trial | | $\mu = 0.005$, $t(15) = 0.120$<br>$P > 0.1$, $d = 0.031$ | $\mu = 0.043$, $t(15) = 0.956$<br>$P > 0.1$, $d = 0.247$ |
| Capture Distance | Recent-2 Trial | $\mu = 0.206$, $t(15) = 4.711$<br>$P < 0.001$***, $d = 1.216$ | $\mu = 0.260$, $t(15) = 6.008$<br>$P < 0.001$***, $d = 1.551$ |
| | Recent-3 Trial | $\mu = 0.202$, $t(15) = 4.405$<br>$P = 0.001$***, $d = 1.137$ | $\mu = 0.253$, $t(15) = 5.877$<br>$P < 0.001$***, $d = 1.518$ |
| ***Hippocampus (Posterior, Right)*** | | | |
| Recent-2 Trial | Capture Distance | $\mu = 0.128$, $t(15) = 2.908$<br>$P = 0.006$**, $d = 0.751$ | $\mu = 0.076$, $t(15) = 1.848$<br>$P = 0.043$*, $d = 0.477$ |
| Recent-3 Trial | | $\mu = 0.112$, $t(15) = 3.418$<br>$P = 0.002$**, $d = 0.883$ | $\mu = 0.047$, $t(15) = 1.233$<br>$P > 0.1$, $d = 0.318$ |
| Capture Distance | Recent-2 Trial | $\mu = 0.241$, $t(15) = 4.700$<br>$P < 0.001$***, $d = 1.214$ | $\mu = 0.222$, $t(15) = 4.951$<br>$P < 0.001$***, $d = 1.278$ |
| | Recent-3 Trial | $\mu = 0.222$, $t(15) = 4.977$<br>$P < 0.001$***, $d = 1.285$ | $\mu = 0.212$, $t(15) = 5.290$<br>$P < 0.001$***, $d = 1.366$ |

**Supplementary Table 5. Functional connectivity of prefrontal and hippocampal clusters**. To examine functional relationships between prefrontal and hippocampal regions, we computed mean (z-scored) functional connectivity values within each cluster based on individual MiOG- and IFG-seed connectivity maps. One-sample permutation *t*-tests (10,000 permutations, one-sided, $P < 0.05$) were applied to assess significance. In both subgroups, the prefrontal clusters exhibited significant and robust cofluctuations. The hippocampal clusters showed stronger connectivity with the MiOG region across both subgroups. μ: mean functional connectivity across participants; *: $P < 0.05$; **: $P < 0.01$; ***: $P < 0.001$.

| Cluster | Subgroup | |
|---|---|---|
| | Discovery | Validation |
| **MiOG-seed FC** | | |
| MiOG *(self)* | $\mu = 0.869$, $t(15) = 21.338$<br>$P < 0.001$***, $d = 5.509$ | $\mu = 0.874$, $t(15) = 29.520$<br>$P < 0.001$***, $d = 7.622$ |
| IFG | $\mu = 0.090$, $t(15) = 3.747$<br>$P = 0.001$**, $d = 0.968$ | $\mu = 0.134$, $t(15) = 5.361$<br>$P < 0.001$**`*, $d = 1.384$ |
| HC (Anterior, L) | $\mu = 0.143$, $t(15) = 7.272$<br>$P < 0.001$***, $d = 1.878$ | $\mu = 0.072$, $t(15) = 2.727$<br>$P = 0.010$**, $d = 0.704$ |
| HC (Anterior, R) | $\mu = 0.134$, $t(15) = 6.176$<br>$P < 0.001$***, $d = 1.595$ | $\mu = 0.064$, $t(15) = 2.831$<br>$P = 0.009$**, $d = 0.731$ |
| HC (Posterior, L) | $\mu = 0.141$, $t(15) = 7.897$<br>$P < 0.001$***, $d = 2.039$ | $\mu = 0.131$, $t(15) = 7.021$<br>$P < 0.001$***, $d = 1.813$ |
| HC (Posterior, R) | $\mu = 0.093$, $t(15) = 4.434$<br>$P < 0.001$***, $d = 1.145$ | $\mu = 0.099$, $t(15) = 5.154$<br>$P < 0.001$***, $d = 1.331$ |
| **IFG-seed FC** | | |
| MiOG | $\mu = 0.066$, $t(15) = 2.801$<br>$P = 0.008$**, $d = 0.723$ | $\mu = 0.127$, $t(15) = 4.711$<br>$P < 0.001$*, $d = 1.216$ |
| IFG *(self)* | $\mu = 0.802$, $t(15) = 23.652$<br>$P < 0.001$***, $d = 6.107$ | $\mu = 0.756$, $t(15) = 20.638$<br>$P < 0.001$***, $d = 5.329$ |
| HC (Anterior, L) | $\mu = 0.048$, $t(15) = 2.084$<br>$P = 0.028$*, $d = 0.538$ | $\mu = 0.008$, $t(15) = 0.675$<br>$P > 0.1$, $d = 0.174$ |
| HC (Anterior, R) | $\mu = 0.046$, $t(15) = 2.047$<br>$P = 0.028$*, $d = 0.529$ | $\mu = 0.003$, $t(15) = 0.123$<br>$P > 0.1$, $d = 0.032$ |
| HC (Posterior, L) | $\mu = 0.039$, $t(15) = 2.554$<br>$P = 0.015$*, $d = 0.659$ | $\mu = 0.005$, $t(15) = 0.247$<br>$P > 0.1$, $d = 0.064$ |
| HC (Posterior, R) | $\mu = 0.028$, $t(15) = 1.420$<br>$P = 0.092$, $d = 0.367$ | $\mu = 0.016$, $t(15) = 0.969$<br>$P > 0.1$, $d = 0.250$ |

### Supplementary Figures

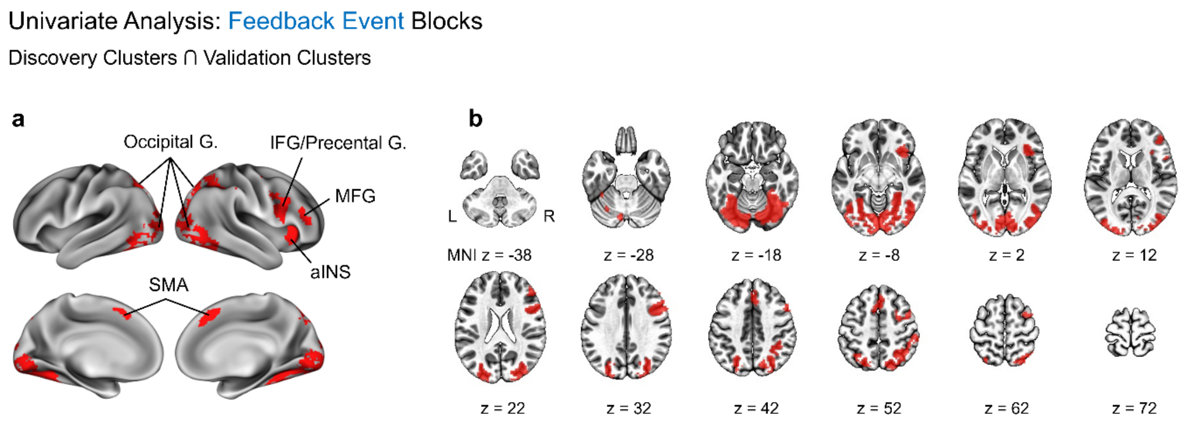


**Supplementary Fig. 1. Univariate analysis of feedback event blocks**. For each subgroup, one-sample *t*-tests were conducted on subject-wise mean beta maps corresponding to all feedback event blocks. Statistical maps were thresholded at voxel-level $P < 0.005$ (one-sided) and cluster-level $\alpha < 0.05$. Brain regions exceeding these thresholds in both subgroups were overlapped, and cross-validated clusters with a minimum extent of 20 voxels were identified.

**a**, Surface-based mapping of the cross-validated clusters. G., gyrus/gyri; IFG, inferior frontal gyrus; MFG, middle frontal gyrus; aINS, anterior insula; SMA, supplementary motor area.

**b**, Volumetric mapping of the identified clusters. No subcortical or cerebellar cluster met the validation criteria across both subgroups.

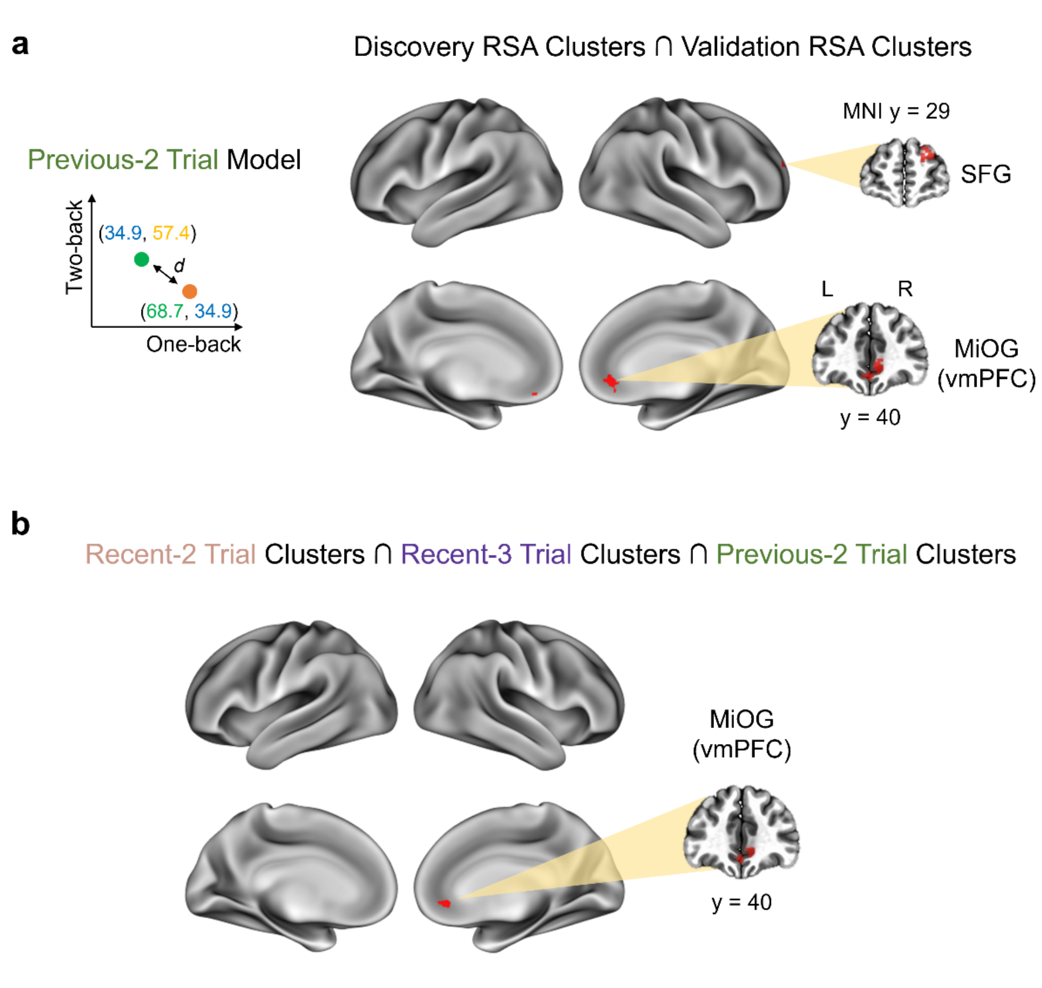


**Supplementary Fig. 2. Cross-validated RSA clusters associated with the Previous-2 Trial model**. This model concatenated one-back and two-back feedback scores for each trial and computed Euclidean distances between feedback score vectors across trial pairs within each run.

**a**, Cross-validated RSA clusters encoding the Previous-2 Trial model. Group-level thresholded inference maps from the two subgroups were overlapped to identify consistent regions. L, left; R, right; SFG, superior frontal gyrus; MiOG, middle orbital gyrus; vmPFC, ventromedial prefrontal cortex.

**b**, A cluster jointly encoding all three multiple-trial feedback history models in both subgroups. The MiOG cluster (24 voxels) appeared in the conjunction map of cross-validated clusters for the Recent-2 Trial, Recent-3 Trial, and Previous-2 Trial models.

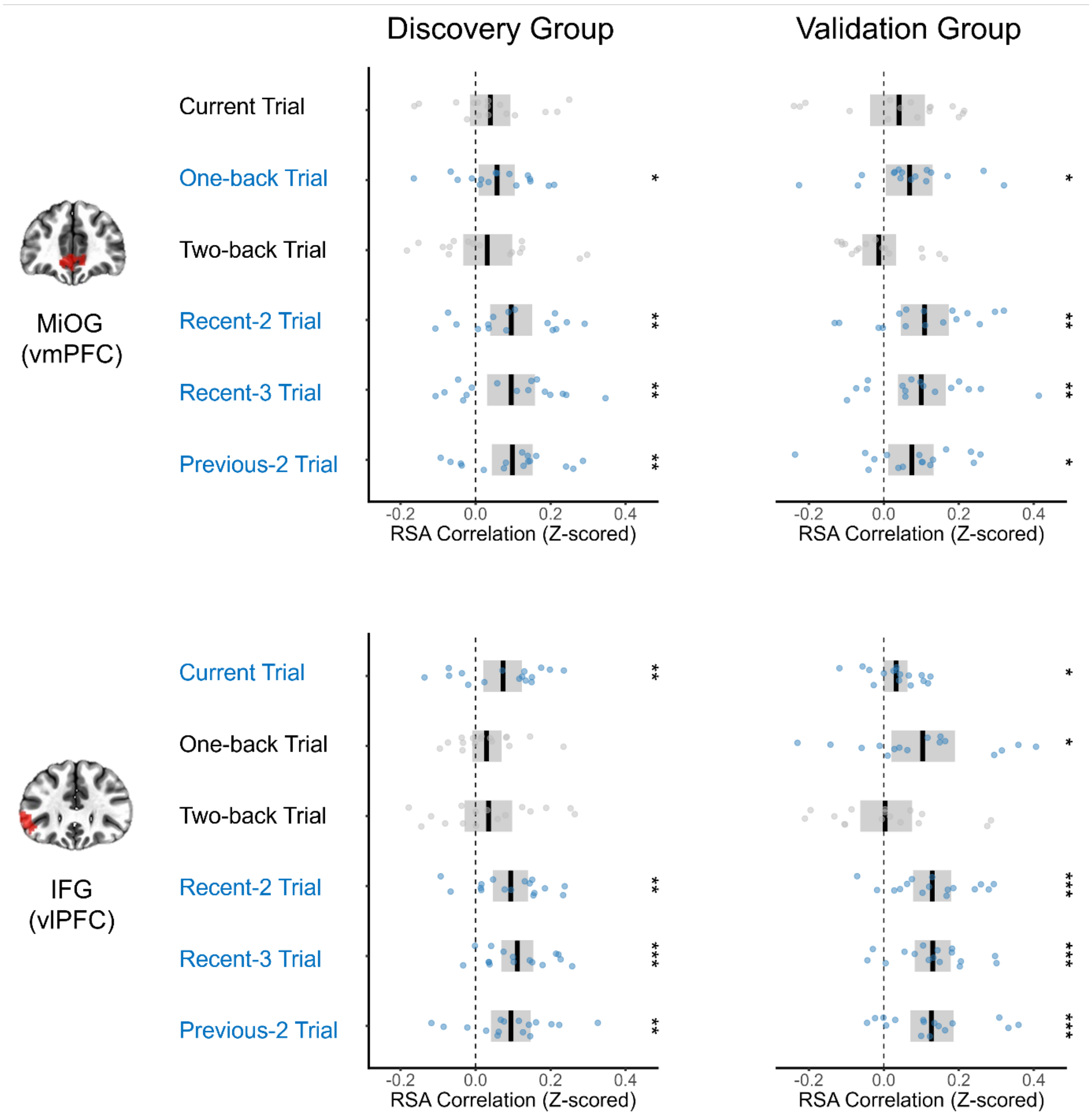


**Supplementary Fig. 3. Second-level RSA of MiOG and IFG clusters with feedback history model RDMs**. Neural RDMs were constructed using all voxels within the MiOG and IFG clusters and compared to all feedback history model RDMs, including control models. One-sample *t*-tests with 10,000 random permutations (null distribution) were conducted for each model. Statistically significant results (one-sided, corrected $P < 0.05$) are highlighted in blue.

Both MiOG and IFG clusters robustly encoded the Recent-2 Trial model (MiOG: Discovery, $t(15) = 3.156$, $P = 0.004$, $d = 0.815$; Validation, $t(15) = 3.207$, $P = 0.003$, $d = 0.828$. IFG: Discovery, $t(15) = 3.805$, $P = 0.001$, $d = 0.982$; Validation, $t(15) = 4.812$, $P < 0.001$, $d = 1.242$); the Recent-3 Trial model (MiOG: Discovery, $t(15) = 2.848$, $P = 0.006$, $d = 0.735$; Validation, $t(15) = 2.949$, $P = 0.004$, $d = 0.761$. IFG: Discovery, $t(15) = 4.954$, $P < 0.001$, $d = 1.279$; Validation, $t(15) = 5.156$, $P < 0.001$, $d = 1.331$); and the Previous-2 Trial model (MiOG: Discovery, $t(15) = 3.387$, $P = 0.002$, $d = 0.875$; Validation, $t(15) = 2.346$, $P = 0.015$, $d = 0.606$. IFG: Discovery, $t(15) = 3.340$, $P = 0.002$, $d = 0.862$; Validation, $t(15) = 4.114$, $P < 0.001$, $d = 1.062$).

Additionally, the MiOG significantly encoded the One-Back Trial model, while the IFG significantly represented the Current Trial model. *: $P < 0.05$; **: $P < 0.01$; ***: $P < 0.001$.

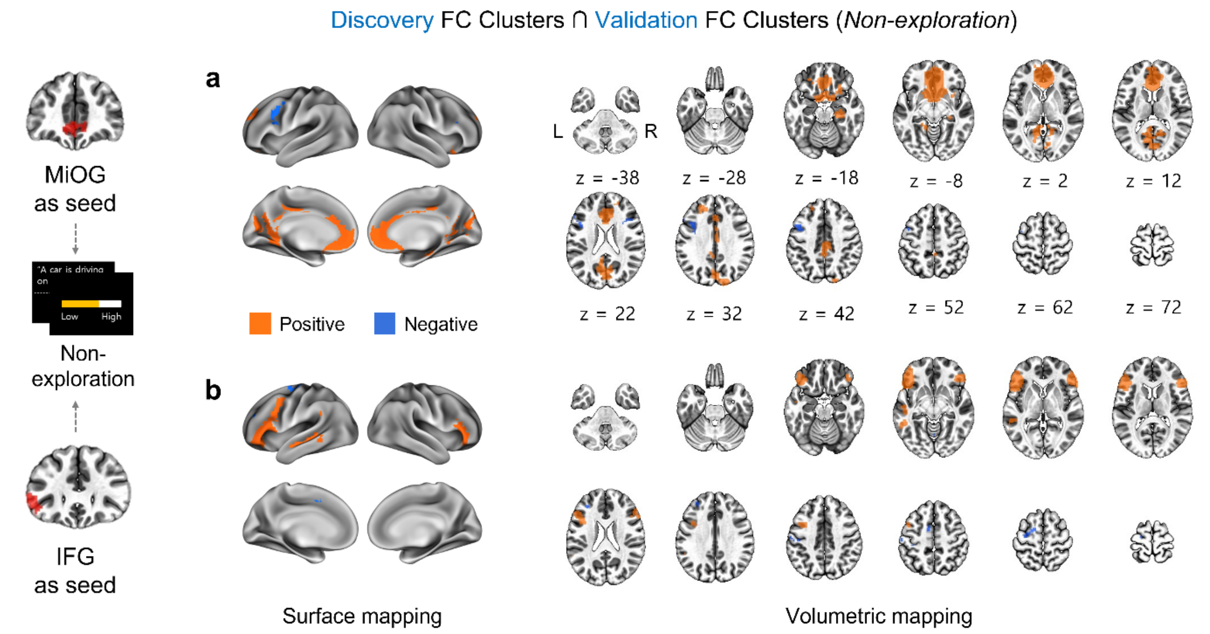


**Supplementary Fig. 4. Cross-validated functional connectivity networks of the MiOG and IFG during nonexploration phases**. We replicated the functional connectivity (FC) analysis pipeline described in Fig. 6, using only nonexploration volumes (i.e., cross-fixations and event blocks outside the street-view environment).

**a**, Surface and volumetric maps of the MiOG-seeded functional network during nonexploration blocks. Orange indicates positive correlations; blue indicates negative correlations.

**b**, Functional network coactivating with IFG during nonexploration.

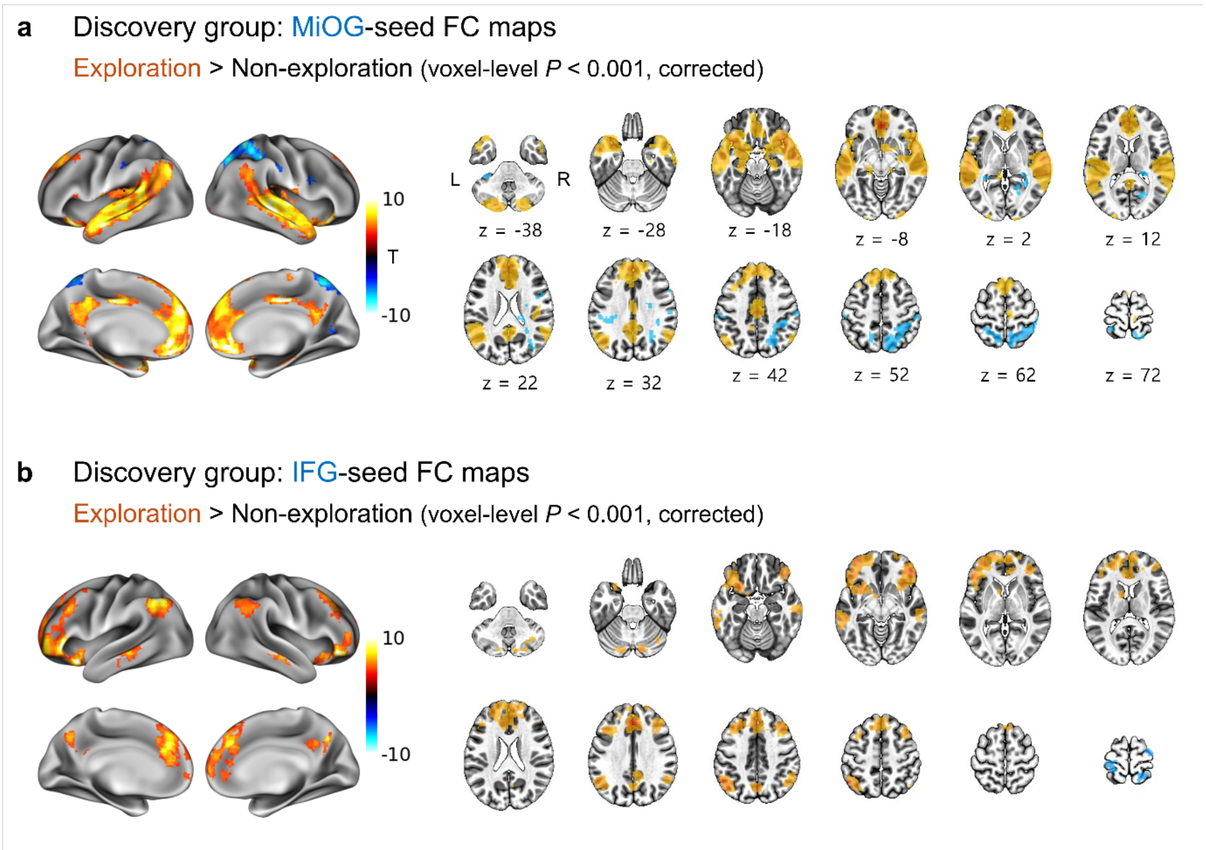


**Supplementary Fig. 5. Differences between exploration and nonexploration functional networks in the discovery group**. Paired *t*-tests were conducted on MiOG- and IFG-seeded functional connectivity maps obtained during exploration phases ("Exploration" networks) and nonexploration blocks ("Nonexploration" networks). Statistical maps were thresholded at voxel-level $P < 0.001$ (two-sided) and cluster-level FDR-corrected $P < 0.05$.

**a**, Brain regions showing significantly greater functional connectivity with the MiOG seed during exploration compared to nonexploration.

**b**, Regions exhibiting stronger functional connectivity with the IFG during exploration phases.

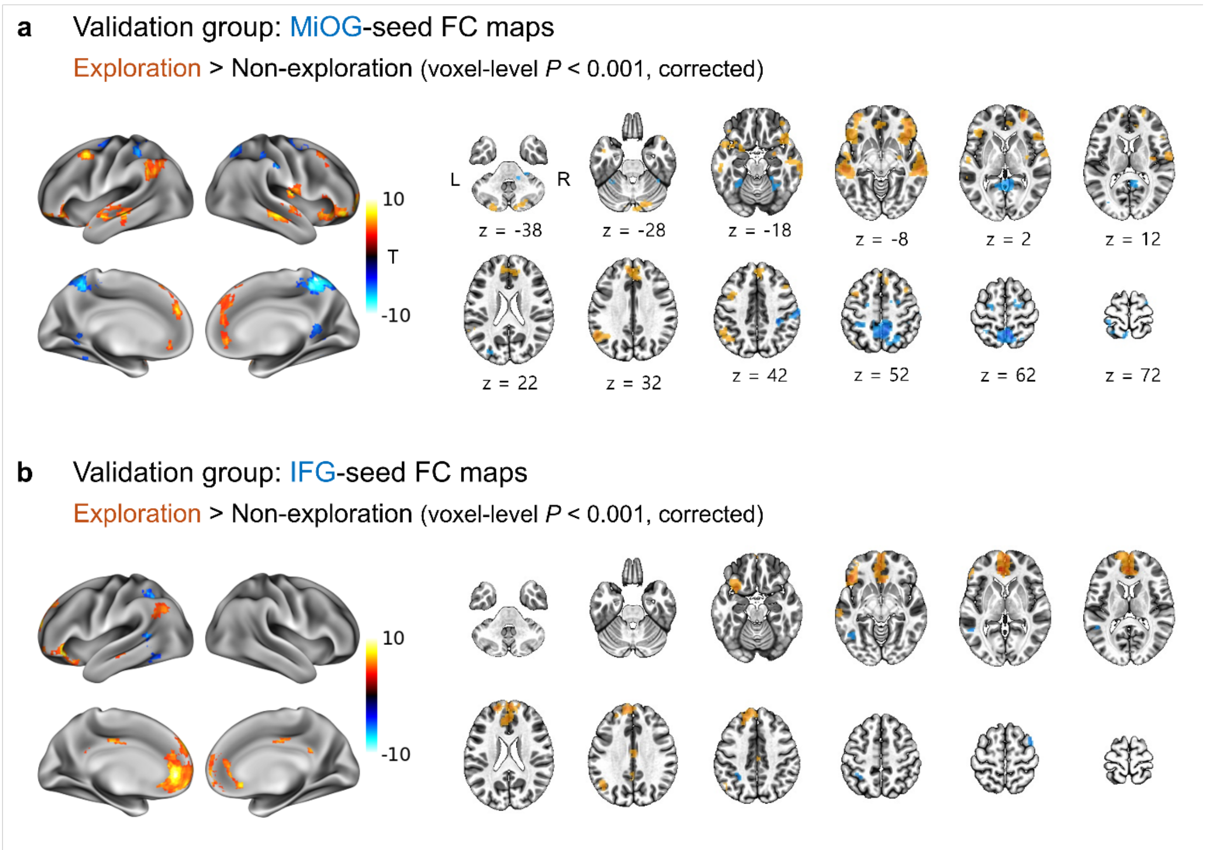


**Supplementary Fig. 6. Differences between exploration and nonexploration functional networks in the validation group**. Paired *t*-tests were repeated in the validation group to compare functional connectivity during exploration and nonexploration phases.

**a**, Clusters showing stronger functional connectivity with the MiOG during exploration blocks.

**b**, Clusters exhibiting significant differences in connectivity with the IFG between exploration and nonexploration phases.

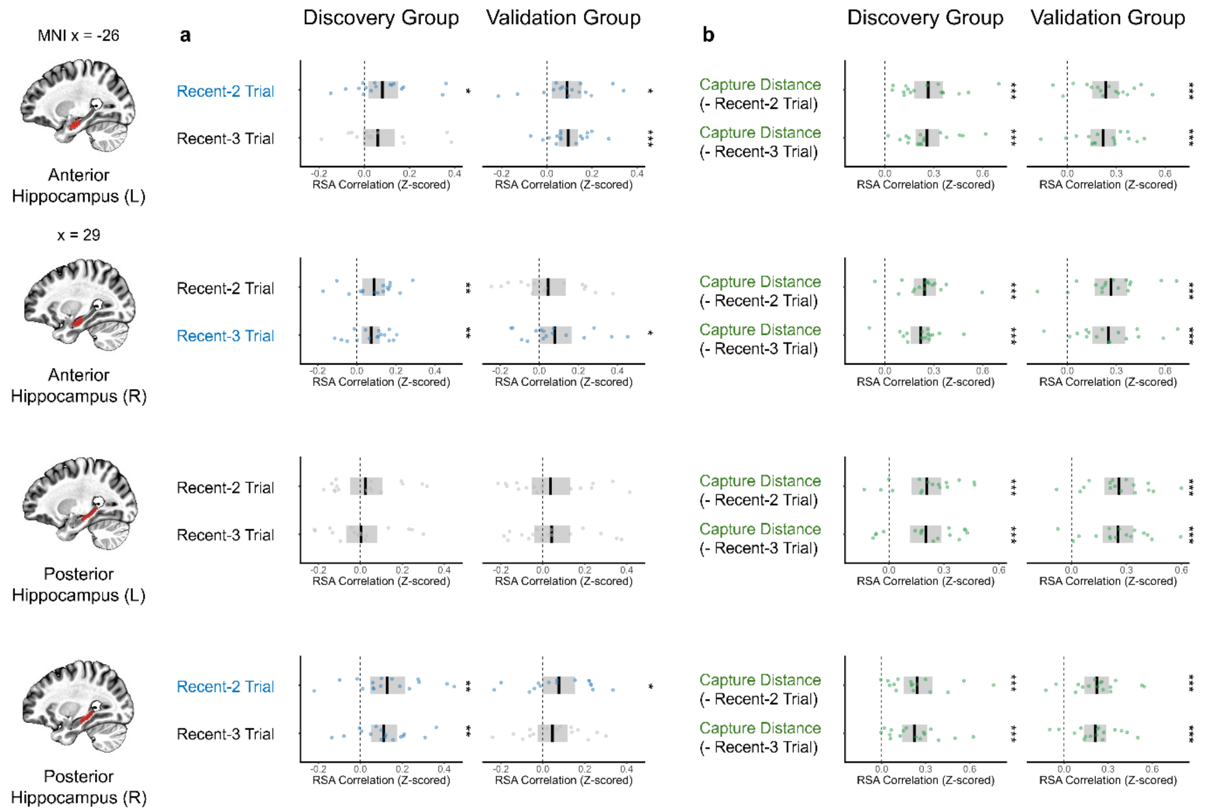


**Supplementary Fig. 7. Partial correlation RSA of hippocampal clusters**. Neural RDMs were constructed using beta values from four hippocampal masks [37], segmented by hemisphere (left/right) and axis (anterior/posterior). These were compared to feedback history model RDMs (Recent-2 Trial and Recent-3 Trial) and the capture distance RDM using a partial correlation approach to minimize confounding between models. L, left; R, right.

**a**, Group-level RSA results for feedback history models after regressing out capture distance effects. Statistically significant associations (one-sided $P < 0.05$, corrected via 10,000 permutations) are shown in blue. The left anterior and right posterior hippocampus significantly encoded the Recent-2 Trial model (left anterior: discovery $t(15) = 2.317$, $P = 0.015$, $d = 0.598$; validation $t(15) = 2.577$, $P = 0.013$, $d = 0.665$; right posterior: discovery $t(15) = 2.908$, $P = 0.006$, $d = 0.751$; validation $t(15) = 1.848$, $P = 0.043$, $d = 0.477$). The right anterior hippocampus encoded the Recent-3 Trial model in both subgroups (discovery $t(15) = 2.983$, $P = 0.006$, $d = 0.770$; validation $t(15) = 1.793$, $P = 0.045$, $d = 0.463$). The left posterior hippocampus did not significantly represent either model.

**b**, Group-level RSA results for the capture distance model after regressing out each feedback history model. All hippocampal regions significantly encoded the capture distance model, even after controlling for the Recent-2 Trial model (discovery: mean $t(15) = 5.498$, all $P < 0.001$, mean $d = 1.420$; validation: mean $t(15) = 5.411$, all $P < 0.001$, mean $d = 1.397$) or the Recent-3 Trial model (discovery: mean $t(15) = 5.714$, all $P < 0.001$, mean $d = 1.475$; validation: mean $t(15) = 5.272$, all $P < 0.001$, mean $d = 1.362$).

*: $P < 0.05$; **: $P < 0.01$; ***: $P < 0.001$.

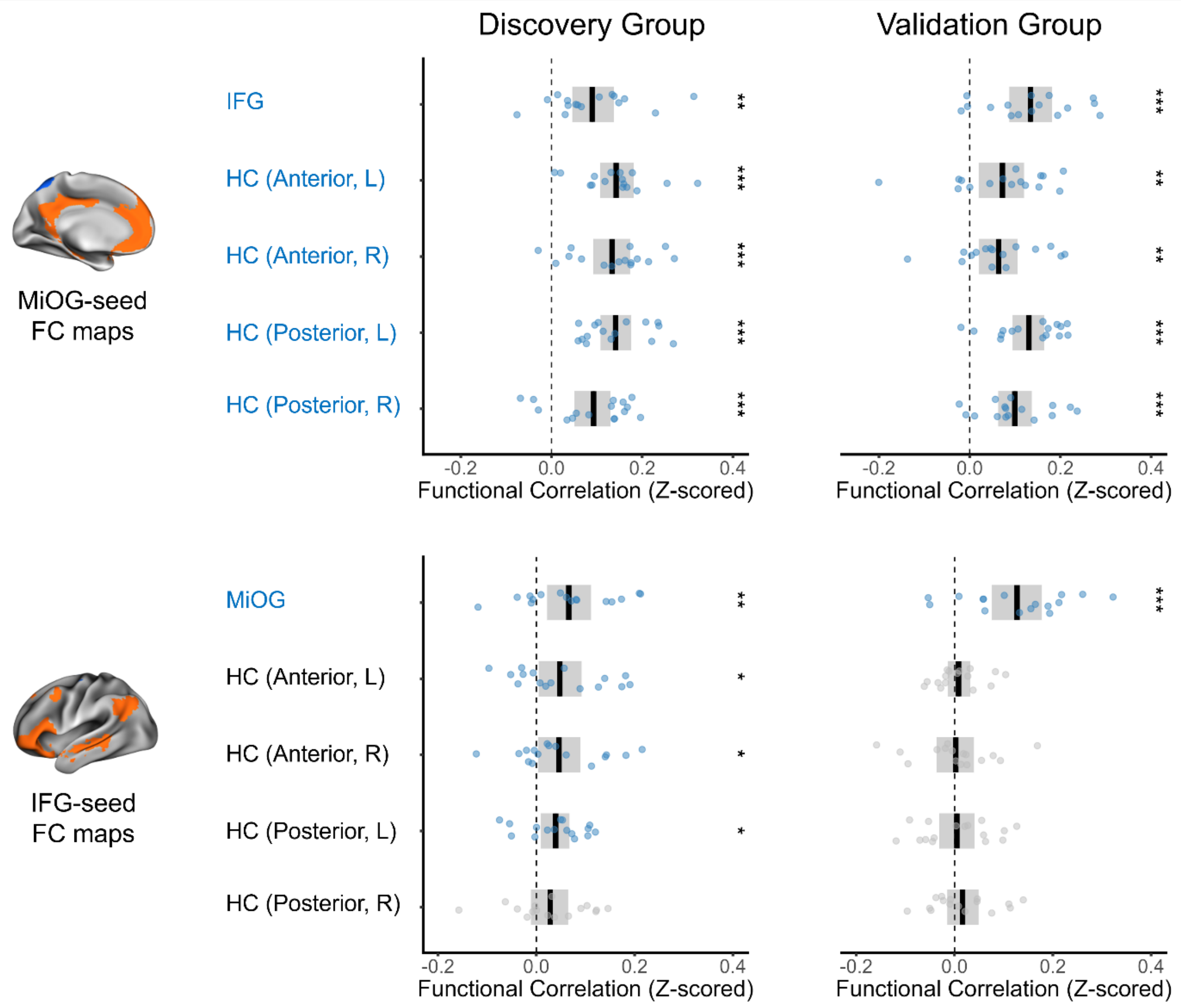


**Supplementary Fig. 8. Mean functional connectivity between MiOG/IFG and hippocampal clusters**. For MiOG- and IFG-seeded functional connectivity (FC) maps, mean *z*-scored correlation values were computed within prefrontal (MiOG and IFG; Fig. 4c) and hippocampal (left/right, anterior/posterior; Supplementary Fig. 7) masks. Permutation-based one-sample *t*-tests (10,000 permutations) were conducted on the individual mean values within each cluster. Cluster names with statistically significant positive associations ($P < 0.05$, one-sided) are highlighted in blue.

In the MiOG-seeded FC maps, all prefrontal and hippocampal clusters showed robust positive connectivity (discovery group: mean $t(15) = 5.905$, all $P < 0.01$, mean $d = 1.525$; validation group: mean $t(15) = 4.619$, all $P < 0.01$, mean $d = 1.193$). In contrast, for the IFG-seeded FC maps, only the MiOG cluster exhibited consistent coactivation across both subgroups (discovery: $t(15) = 2.801$, $P = 0.008$, $d = 0.723$; validation: $t(15) = 4.711$, $P < 0.001$, $d = 1.216$). Additionally, some hippocampal clusters in the discovery group (left/right anterior hippocampus and left posterior hippocampus) showed significant coactivation (mean $t(15) = 2.228$, all $P < 0.05$, mean $d = 0.575$).

HC, hippocampus; L, left; R, right; *: $P < 0.05$; **: $P < 0.01$; ***: $P < 0.001$.

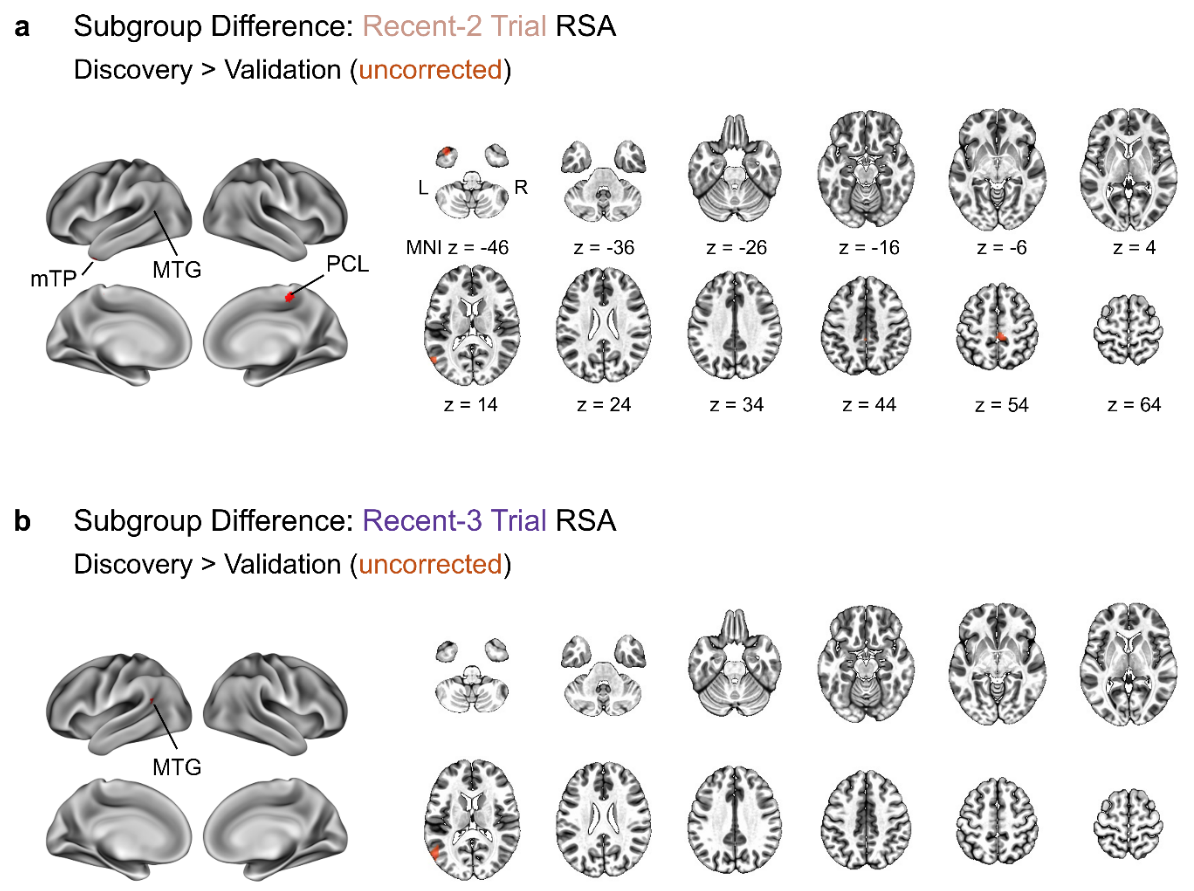


**Supplementary Fig. 9. Subgroup differences in the representational strength of feedback history models**. Two-sample independent *t*-tests were conducted on individual feedback history RSA maps (averaged across runs) from both subgroups using AFNI's 3dttest++. Clusters with stronger representational strength in the discovery group than in the validation group were identified at a one-sided voxel-level threshold of $P < 0.005$ and a minimum cluster size of 20 voxels. However, no clusters survived cluster-level correction using 3dClustSim. Additionally, no clusters showed greater representational strength in the validation group compared to the discovery group at the same threshold.

**a**, Uncorrected clusters where the Recent-2 Trial model showed stronger encoding in the discovery group. L, left; R, right; mTP, medial temporal pole; MTG, middle temporal gyrus; PCL, paracentral lobule.

**b**, An uncorrected cluster (MTG) with higher representation of the Recent-3 Trial model in the discovery group.

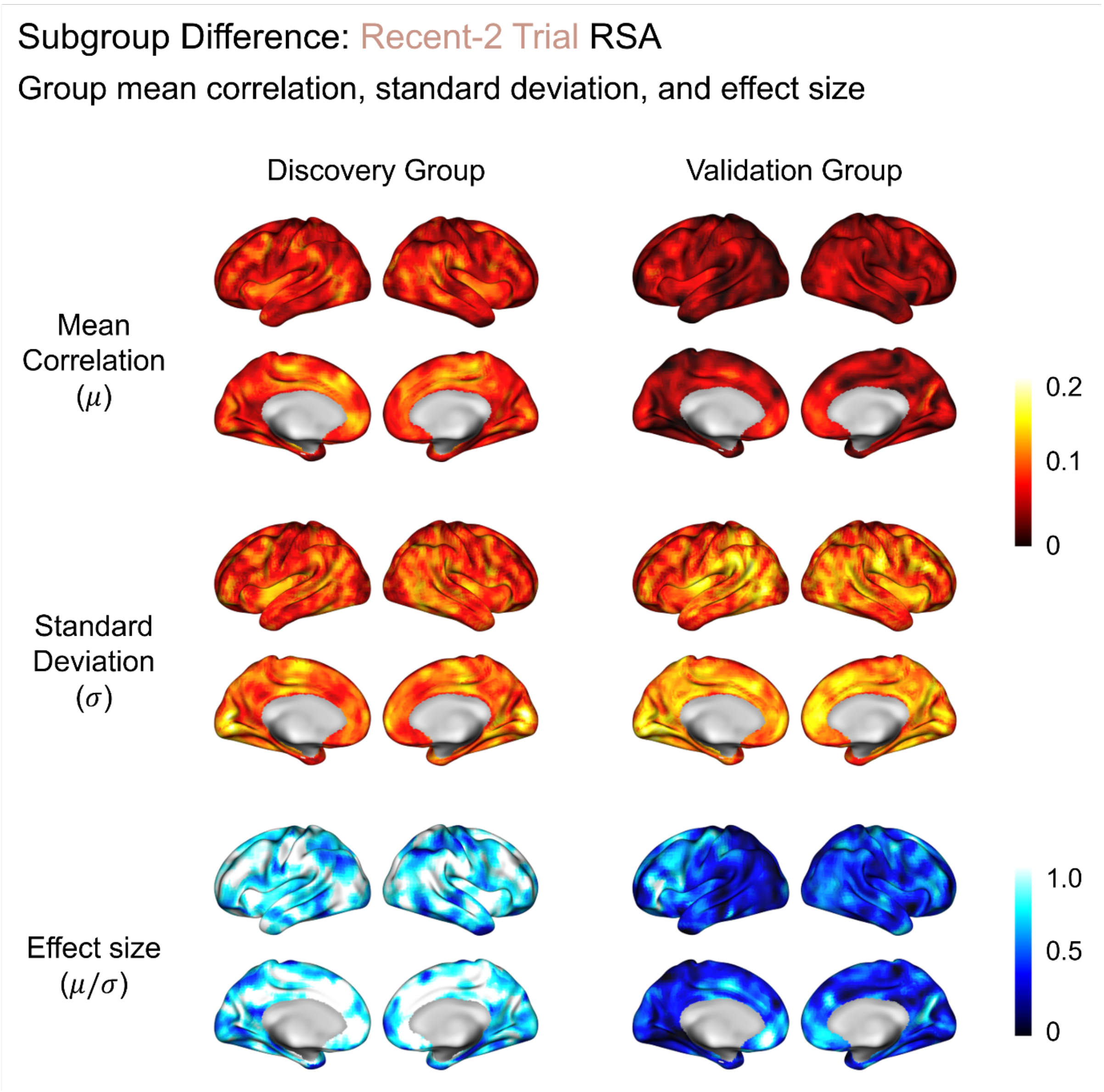


**Supplementary Fig. 10. Group mean correlation, standard deviation, and effect size for the Recent-2 Trial RSA**. For each subgroup, the mean correlation (μ) from individual Recent-2 Trial RSA maps, the standard deviation (σ), and the standardized effect size ($d = \mu/\sigma$) were computed and visualized on the fsLR template. Notably, the validation group exhibited greater voxelwise variability, particularly in the parietal, temporal, and occipital regions, resulting in lower effect size estimates compared to the discovery group.

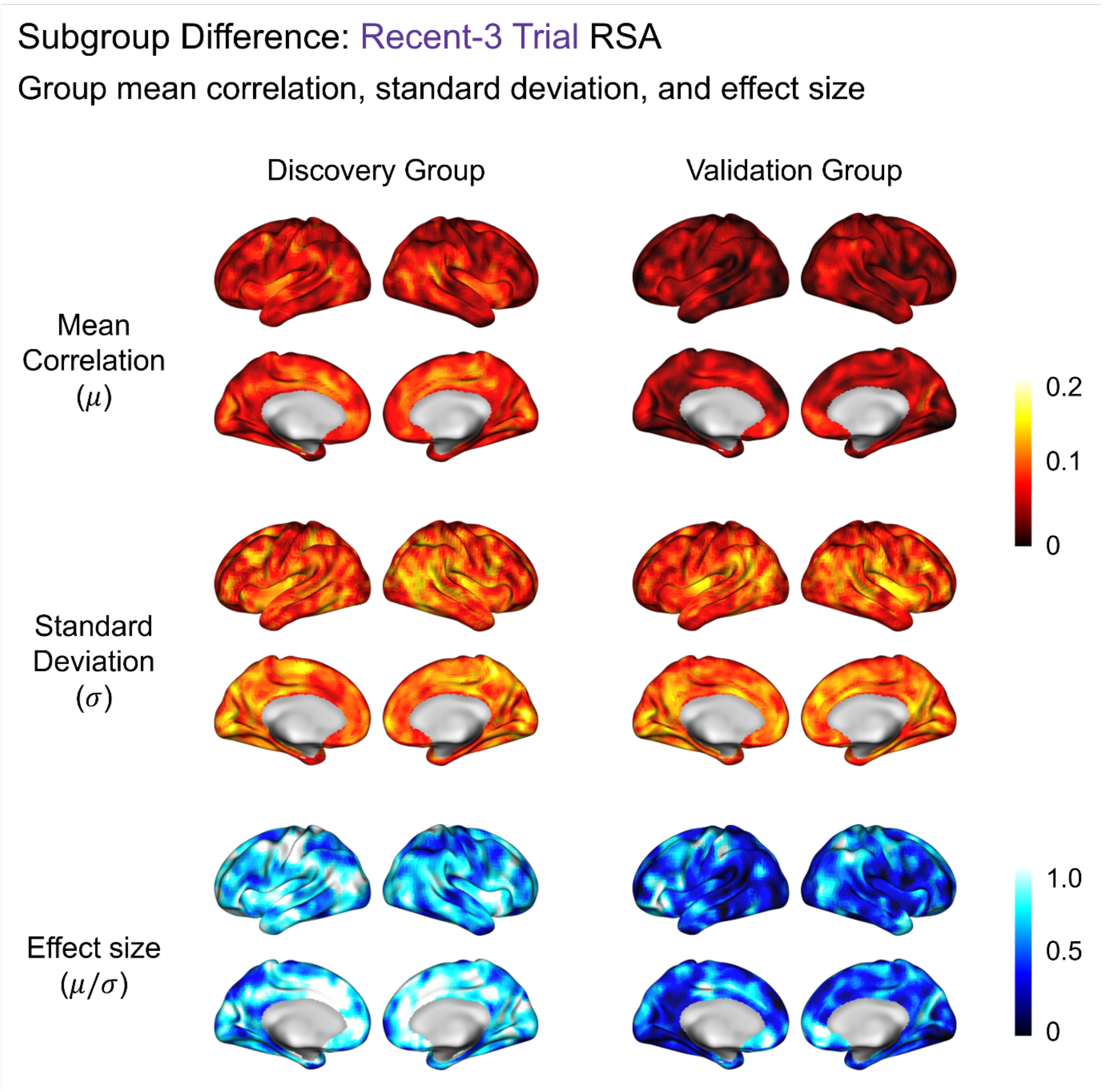


**Supplementary Fig. 11. Group mean correlation, standard deviation, and effect size for the Recent-3 Trial RSA**. Voxelwise mean correlations (μ), standard deviations (σ), and effect sizes ($d$ = μ/σ) for the Recent-3 Trial model are presented. Unlike the Recent-2 Trial model, standard deviation estimates for the Recent-3 Trial model are comparable between the two subgroups.
